# Accelerating discovery and integration from RNA to Protein with *R2T2P*

**DOI:** 10.64898/2026.11.05.764226

**Authors:** Roberto Albanese, Amy E. Campbell, Matteo Bonfanti, Azmal Syed Ali, Chela T. James, Andrea Graziadei, Giorgio Tasca, Alberto Riva, Jeroen Krijgsveld, Sujatha Jagannathan, Lorenzo Calviello

**Affiliations:** Human Technopole, Milan, Italy; Department of Biochemistry and Molecular Genetics, University of Colorado Anschutz Medical Campus, Aurora, CO 80045, USA; Division of Proteomics of Stem Cell and Cancer, German Cancer Research Center (DKFZ), Heidelberg, Germany; Medical Faculty, Heidelberg University, Heidelberg, Germany; John Walton Muscular Dystrophy Research Centre, Newcastle University; The Newcastle upon Tyne Hospitals NHS Foundation Trust; and NIHR Biomedical Research Centre (BRC): Newcastle, Newcastle upon Tyne, United Kingdom; RNA Bioscience Initiative, University of Colorado Anschutz Medical Campus, Aurora, CO 80045, USA

## Abstract

Integrating transcriptomics and proteomics data remains an unmet challenge in the post-genomics era, discouraging researchers from pursuing integrative approaches to molecular biology research. Leveraging the power of translation profiling, we present RNA to Translation to Protein (R2T2P), a computational platform integrating genome-guided transcriptome assembly with identification of translation and proteins, including a tailored differential analysis framework and support for multiple reference transcriptomes. We applied R2T2P to investigate a cellular model of facioscapulohumeral muscular dystrophy (FSHD), where ectopic expression of the *DUX4* retrogene in skeletal muscle triggers significant alterations in transcription and RNA metabolism. We uncovered hundreds of uncharacterized, highly overexpressed RNAs, including antisense and alternative transcripts, and quantified their contribution to translatome and proteome. Re-analysis of expression datasets from patient-derived myotubes and muscle biopsies validated this alternative transcriptome in FSHD patients, adding a reservoir of *bona-fide* novel molecular actors in a rare disease. Available at https://github.com/nfdata-omics/R2T2P/, R2T2P is a flexible and powerful workflow in modern integrative biology.

## Introduction

With incredible advances in high-resolution monitoring of both RNA and protein species, a fundamental challenge of current biomedical research lies in the joint analysis of this wealth of information to develop an integrative view of the molecular complexity of the cell^1^. Large differences in the biochemistry of nucleic acids and proteins, as well as technologies and data analysis solutions devoted to their quantification, greatly complicate this task, acting as significant obstacles towards more comprehensive, multimodal approaches to molecular biology research^2,3^. Moreover, a one-gene-one-protein framework is often applied when addressing these integrative efforts, ignoring the wide functional diversity of the molecular species involved, and potentially misrepresenting molecular mechanisms^4^.

With more than 200 thousand annotated transcripts^5^ and a similar number of expressed RNAs obtained via a large collection of long-read sequencing data^6^, it is an arduous task to determine their contribution to the proteome. This task is complicated by alternative transcript isoforms for most protein-coding genes, many of which can produce protein isoforms or have unclear coding potential. Moreover, thousands of abundant long non-coding RNAs (lncRNAs), transcribed from intergenic space or antisense to known genes, emerged as a new class of regulators, often via direct interaction with chromatin^7^, or sometimes as templates for protein synthesis^8^.

The expression of numerous RNAs can be confined to a specific cell type or state, or exhibit disease specificity^9^. While significant efforts are devoted to providing high-quality gene models, novel RNAs are routinely being discovered in a variety of tissues, cell states, and diseases^10,11^. However, their regulation, as well as their contribution to the translated transcriptome, is often overlooked.

Translation profiling (in particular, the Ribo-seq protocol^12^) offers a powerful solution to identify the ensemble of translated regions^13^, and quantify the contribution of complex transcriptomes to protein synthesis^14^. While increasing attention is being given to the detection of translation of small open reading frames^15^, little effort has been directed towards the unbiased quantification of translation in complex transcriptomes and isoform-level regulation, while accounting for the presence of novel translated loci.

Modern advances in mass spectrometry workflows and instrumentation have substantially increased proteome coverage^16^, enabling deeper investigation of proteins encoded by alternative transcript isoforms^17^, and calling for the development of integrative approaches between proteomics and nucleotide-based -omics methods^18^. However, existing proteogenomic workflows often focus on the detection of specific types of proteins, such as peptides coming from translation of small ORFs^19,20^, N-terminal proteoforms^21,22^, potentially immunogenic peptides^23–25^, as well as peptides produced as a consequence of DNA mutations^26^ or alternative RNA isoforms^27,28^. Importantly, the presence of multiple, often unannotated transcripts that could give rise to novel peptides greatly complicates interpretation, and can lead to ambiguous identification of non-canonical translation mechanisms^29,30^.

Given the importance of -omics profiling in human disease, integrative and discovery-oriented approaches will advance our understanding of disease mechanisms, particularly in conditions characterized by markedly altered post-transcriptional processes. One example is facioscapulohumeral muscular dystrophy (FSHD), one of the most prevalent genetic muscle diseases, linked to aberrant expression of *DUX4* ^31–33^. *DUX4* encodes an embryonic transcription factor normally silenced in skeletal muscle, which causes a dramatic change in gene expression and RNA metabolism in cellular models mimicking DUX4 overexpression^34–36^. If such alterations create an aberrant cellular proteome, and how this leads to a progressive loss of muscle functions, is currently unknown. More broadly, a comprehensive, isoform-aware workflow able to quantify gene expression (mis)regulation from RNA all the way to translation and protein is currently lacking.

To this end, we here present R2T2P (*RNA to Translation to Protein*), a flexible, integrative workflow linking identification of RNAs with active translation and proteome-wide quantification, including tailored quality control and isoform-aware differential analysis solutions. Applied to a comprehensive dataset from cellular models, R2T2P quantifies molecular novelty at different layers of the gene expression cascade, and enables re-analysis of patient expression data to uncover previously unrecognized disease biology.

## Results

### The R2T2P workflow

To link discovery and quantification across different assays, we organized *R2T2P* into three interconnected modules (**Figure 1**), defined as follows (**Methods**). The RNA module uses RNA-seq data, as well as genome and reference .gtf file including CDS annotation (e.g. from GENCODE), to find novel junctions with *STAR*^37^ (**Methods**), followed by merging of detected junctions and performing genome-guided transcriptome assembly with *StringTie3*^38^, in conservative mode. This new annotation is compared and merged with the reference annotation, using *gffcompare*^39^; in this step, an additional *.gtf* file (e.g. derived by long-read assemblies) can also be included, and all transcript structures are merged and harmonized (**Methods**) in the final annotation, tracing the annotation source for each gene and transcript feature. All computation on genomic and transcriptomics features is performed using the power, efficiency and flexibility of different packages in the Bioconductor^40,41^ project, such as *GenomicFeatures*^42^, *rtracklayer*^43^, and many others (**Methods**). By expanding the reference transcriptome, this first module establishes the foundation for tracking the expression of known and novel RNAs throughout all subsequent steps of the pipeline.

**Figure 1:**
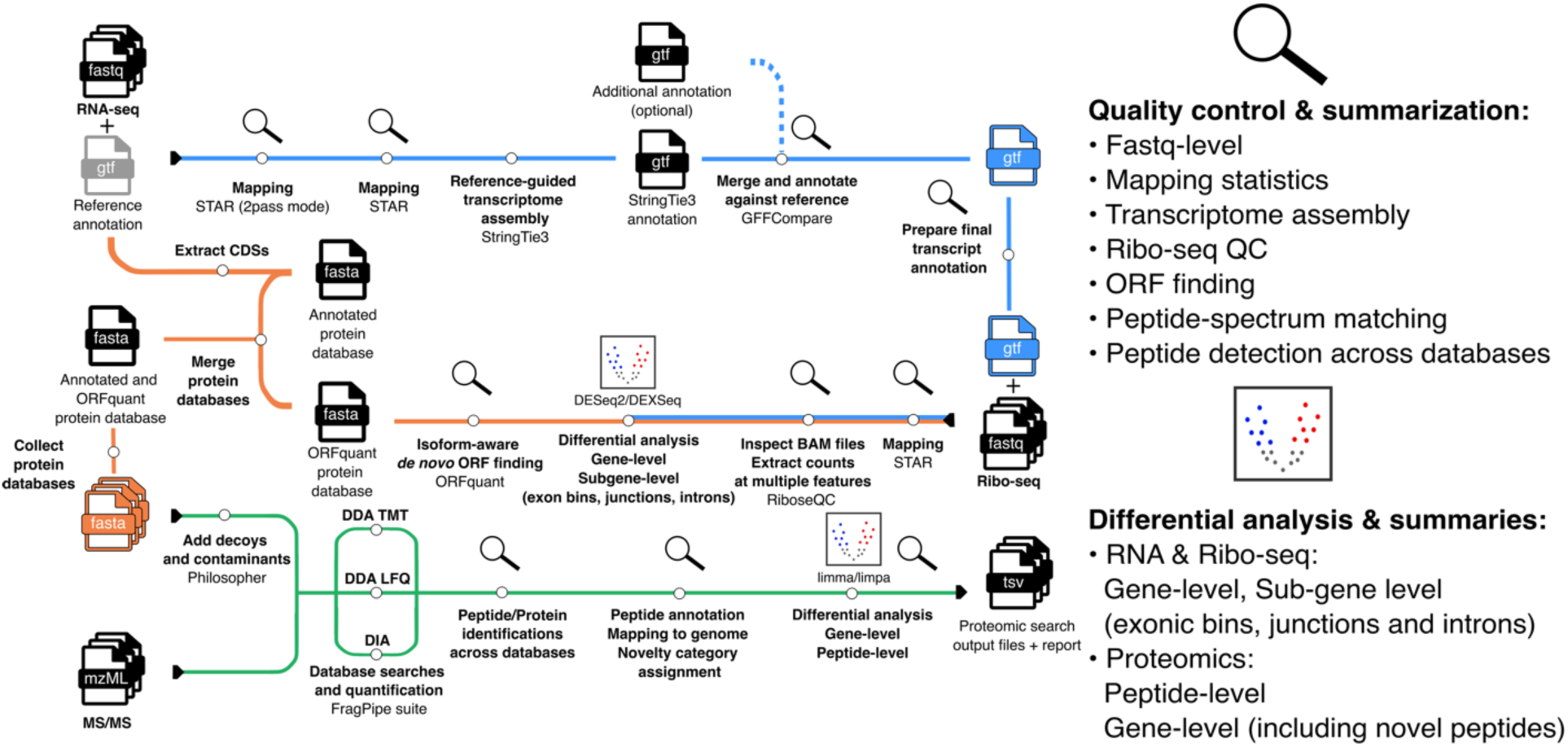
The R2T2P workflow. A schematic of the entire pipeline; main steps are shown in bold, with each module (RNA, Translation, Protein) indicated in blue, orange and green, respectively; software tools and file formats are indicated, as well as QC (magnifying glasses) and differential analysis steps (volcano plots).

In the Translation module Ribo-seq data is added and mapped (disabling novel junction discovery) using the final annotation; quality control of the resulting mapping and extraction of counts and other statistics is performed with *RiboseQC*^44^, allowing for the first set of differential analyses: count matrices on coding sequences, exons, and flattened gene models (exonic bins, junctions and introns) are used to perform differential analysis at the gene level, using counts on exons or CDS with *DESeq2*^45^, as well as at the sub-gene level, using counts on flattened gene models with *DEXSeq*^46^. For both RNA-seq and Ribo-seq, count matrices are computed using all reads or uniquely mapping reads, enabling analyses focused on unambiguously mappable regions. In addition, R2T2P includes extra differential analysis steps, as well as post-processing, described in the (**Methods**) section.

Finally, P-sites positions are automatically extracted using *RiboseQC*^44^ from Ribo-seq data, as input for *ORFquant*^14^, which performs isoform-aware translation detection and quantification, allowing to monitor translation on novel RNAs. This step generates a list of translated ORFs and a custom protein database to be included in the next steps of the workflow.

In the Protein module, ORFquant-derived protein sequences, annotated protein sequences, and their combination are used to generate three different databases in .*fasta* format. Contaminants and decoys are then added to the databases with *Philosopher*^47^. The integrative *FragPipe* suite^48–50^ is then used to perform peptide database searches and quantification. According to the type of proteomics data, different workflows (DIA, DDA-LFQ, DDA-TMT) are used, each comprising different software tool combinations for identification, re-scoring, quantification, and other post-processing steps (**Methods**). The threshold for false-discovery-rate (FDR) is set to 0.01 at PSM, and peptide, and protein-level, without performing separate FDR control for novel peptides (**Discussion**). Differential analysis is performed at peptide-and gene-level with *limpa*^51^, a dedicated *limma*^52^-based package for differential analysis of proteomics data including model-based imputation solutions, using either raw intensities or *limpa*-imputed values.

Multiple QC checks are included at each step, including associated visualizations of obtained results for both identification, and differential analysis. The entire R2T2P workflow is implemented in Nextflow^53^, improving workflow standardization, reproducibility, and portability across different computing environments. For its implementation, we used the nf-core framework and its associated tooling, including the nf-core pipeline template, tools, and modules^54^. Additionally, runtime and other statistics on compute resource usage at each individual step are provided (**Supplementary Figure 1**). More details on individual R2T2P steps and output are available in the **Methods** sections.

### Hundreds of novel DUX4 targets in human muscle cells

We applied the full R2T2P workflow to a time series of a human muscle cell line (MB135) following ectopic induction of *DUX4* ^55^, which we refer to as “iDUX4 cells”. The time series comprises RNA-seq, Ribo-seq^34^, and TMT-labeled mass spectrometry data, performed in triplicate (**Figure 2A**, **Methods**). Despite modest sequencing depth (**Supplementary Table 1**), we identified numerous novel transcripts, of which 7591 are alternative isoforms of known coding genes, and 1662 contribute to annotation for 1532 novel genes. Novel isoforms of coding genes displayed the expected length distribution, while transcripts of novel genes were much shorter (**Supplementary Figure 2**). Focusing on the gene level, 361 novel genes belonged to the antisense category, 570 mapped antisense to introns of known genes, and 601 to intergenic loci (**Figure 2B**).

**Figure 2:**
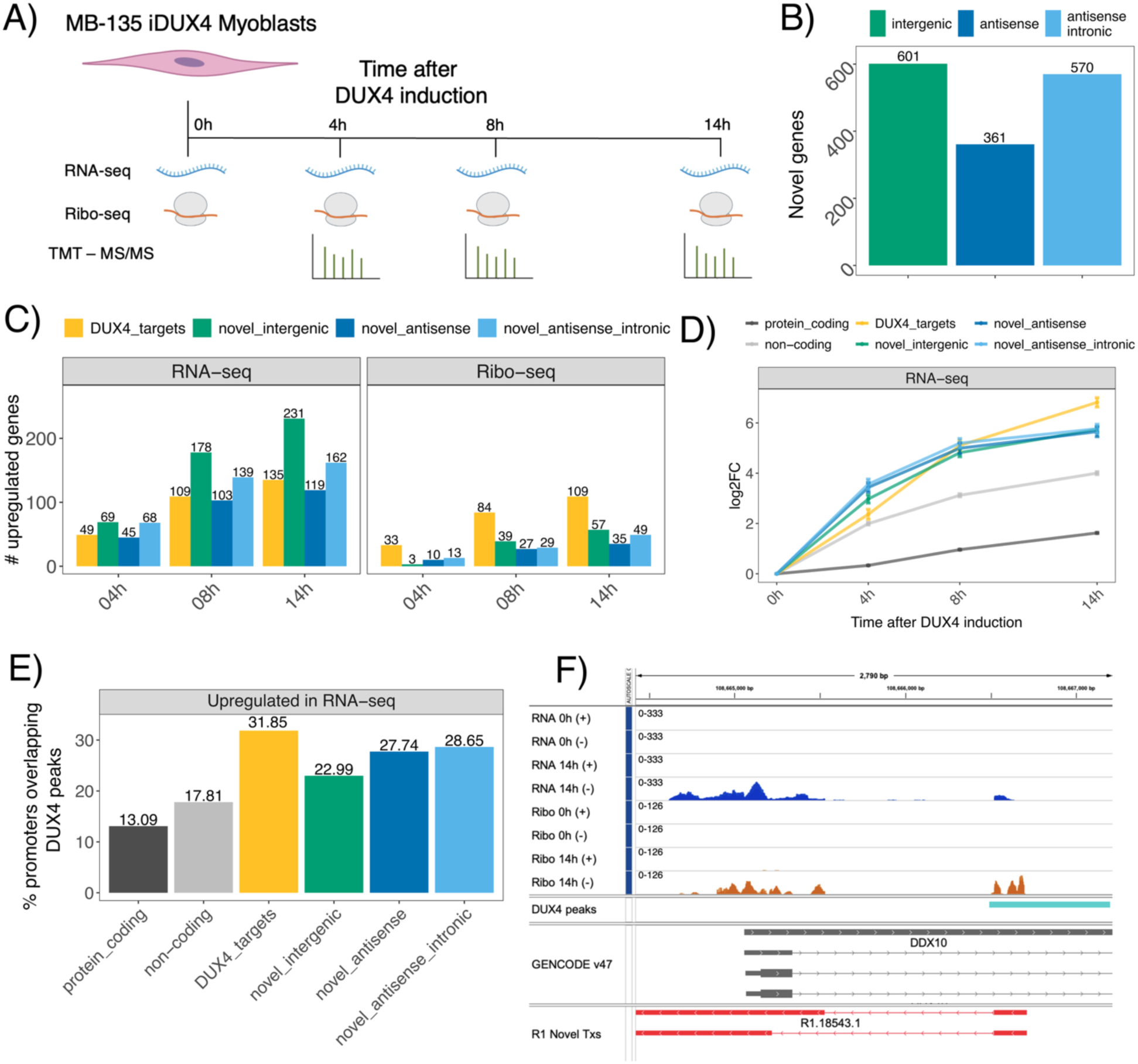
Novel DUX4 target RNAs in human muscle cells. A) schematics of the human muscle cells (iDUX4) experimental setup. B) Number of novel genes detected by *StringTie3*. C) the number of upregulated genes across different annotation categories, including known DUX4 targets (in gold), at different time points after DUX4 induction. D) the Log2FC (y-axis) of the different categories over time (x-axis), showing similar dynamics between novel genes and known DUX4 targets. E) the overlap between annotated ChIP-seq peaks and promoters from different gene categories. F) an example showing the *DDX10* locus: RNA-seq and Ribo-seq tracks at 0h and 14h post-DUX4 induction are shown above, with DUX4 peaks, GENCODE v47 annotation and *StringTie3*-derived novel transcripts below; of note, the annotated *DDX10* gene is on the “+” strand, with the novel RNA on the “-” strand.

Using a list of consensus genes regulated by DUX4 (n=183 mappable to GENCODE v47 ids, which we will refer to as “DUX4 targets”) as a positive control^32^, we quantified the contribution of novel genes to the DUX4-regulated transcriptome. After filtering for excessively overlapping loci (**Methods**) we detected numerous upregulated genes across the time course (**Supplementary Figure 3, Supplementary Table 2**), including 512 novel genes at 14h post-DUX4 induction (**Figure 2C, Methods**), compared to 135 known DUX4 targets. When using Ribo-seq data, we detected fewer overall upregulated genes, especially novel genes, possibly due to their noisier expression levels as well as to their potentially non-coding status.

Both upregulated known and novel genes displayed a similar level of induction at different timepoints after *DUX4* activation (**Figure 2D**), including in Ribo-seq (**Supplementary Figure 4**), raising the hypothesis that they represent additional direct DUX4 targets. To test this, we quantified the overlap of annotated DUX4 ChIP-seq peaks^56^ with promoters of known targets and novel genes (**Methods**). As shown in **Figure 2E**, promoters of novel genes (for all the novel categories) were enriched for DUX4 binding compared to other upregulated genes. Such enrichment was stronger when using genes that were upregulated in Ribo-seq data (**Supplementary Figure 5**). One example is shown at the *DDX10* locus (**Figure 2F**), where a novel antisense RNA is markedly upregulated in both RNA-seq and Ribo-seq, reaching expression levels well above the annotated locus.

Enabled by the capabilities of the *R2T2P* workflow, we expanded the set of DUX4 target genes, including hundreds of novel RNAs otherwise invisible with standard analytical pipelines.

### An expanded view on DUX4-induced protein synthesis

Leveraging the high quality of our Ribo-seq data^34^ and the power of *R2T2P* Translation and Protein modules, we quantified the contribution of the DUX4-induced transcriptome to protein synthesis. Using uniquely mapping Ribo-seq reads, we detected and quantified translation of 26113 ORFs in 13923 genes across a variety of different biotypes (**Supplementary Figure 6**, **Supplementary Table 3**). Thanks to our augmented transcriptome, we detected 1396 translated ORFs uniquely originating from novel transcripts (**Figure 3A**). Of these, 1151 mapped to novel isoforms of known genes and 245 to transcripts in novel genes, with lengths and expression levels similar to annotated ORFs and ORFs in non-coding RNAs, respectively (**Supplementary Figure 6**). When using all Ribo-seq reads numbers were similar, except for detected ORFs on pseudogenes (**Supplementary Figure 7**).

**Figure 3:**
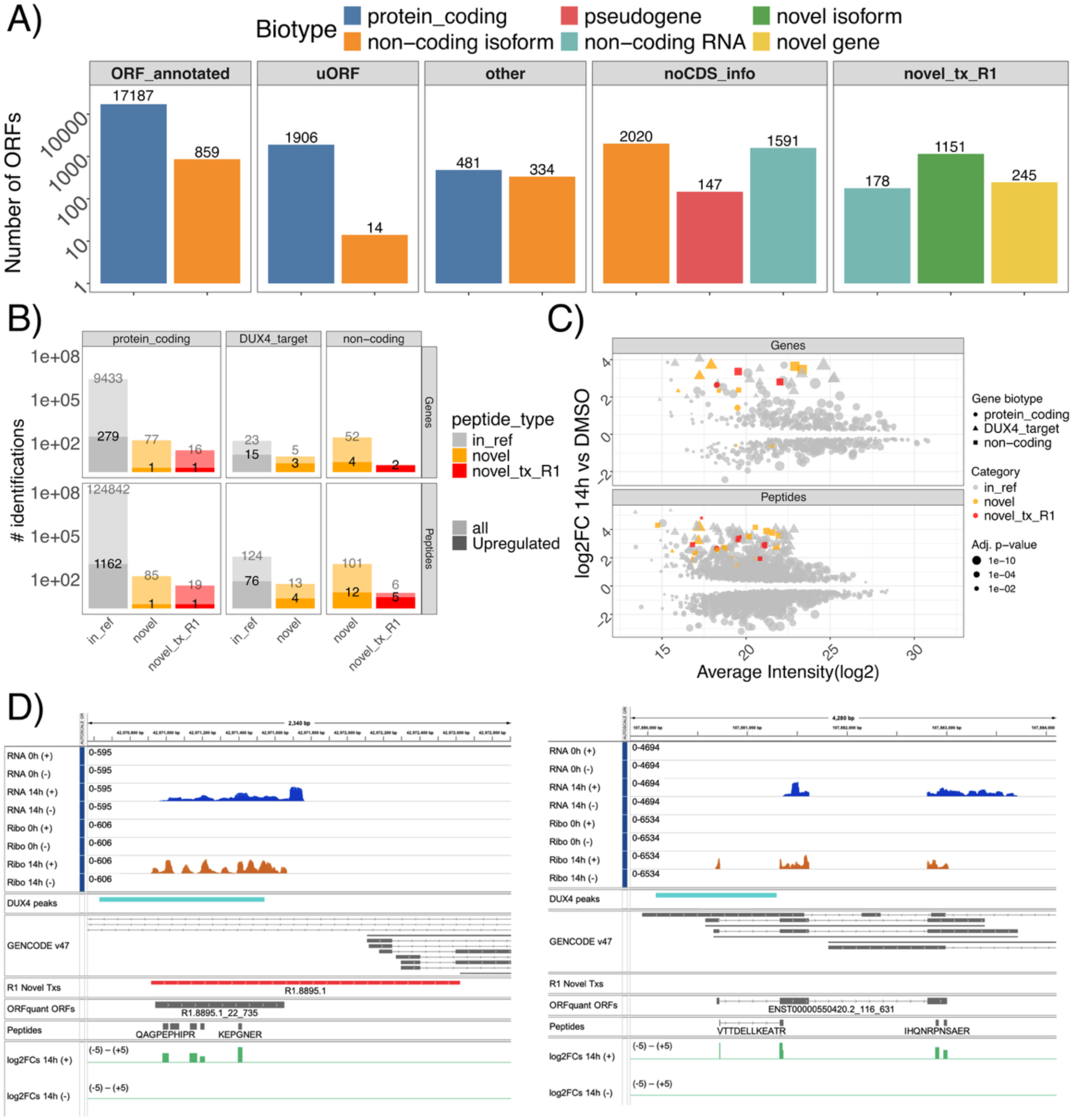
Unbiased identification of DUX4-induced translation and protein products. In A) Number of quantified translated ORFs by *ORFquant* across different biotypes. Number of detected peptides and their corresponding genes in B), with numbers of significantly upregulated species also specified in darker colors. In C) an MA plot with peptide-level abundance (x-axis) against log2FC at 14h post DUX4 induction (y axis). In D) two examples showing a novel RNA (on the left), and a non-coding locus (on the right); same tracks as in Figure 2F, adding *ORFquant*-detected ORFs, detected peptides, and peptide-level log2FC values.

We then assessed the ability of this augmented map of mRNA translation as an enhanced protein database to quantify protein abundance and regulation across DUX4 induction, using the R2T2P Protein module supplied with deep TMT-labeled mass spectrometry data (**Methods**). Combining the *ORFquant*-derived protein database and all annotated ORFs in GENCODE, we detected 134619 peptides, of which 125190 mapped to one gene, providing quantification for 9517 genes (**Figure 3B**, **Supplementary Table 4**). In addition, 224 peptides mapped to ORFquant entries only, including 25 mapping to ORFs in novel transcripts. Most peptides were commonly detected by the single database searches alone (**Supplementary Figure 8**); while some peptides were identified by the ORFquant database only (**Discussion**), we decided to focus on results from the combined database search. At 14 hours post-DUX4 induction, 1261 peptides in 456 genes were found to be significantly upregulated, with few upregulated peptides coming from novel ORFs, or novel ORFs from novel transcripts (**Figure 3B**). Despite the low number of identifications, upregulation levels placed these novel species among the most regulated proteins upon DUX4 induction, at levels comparable to well-known DUX4 targets (**Figure 3C**). **Figure 3D** shows the two most striking examples: one comprising 5 peptides from a novel ORF on a novel transcript, and one with 4 peptides from a novel ORF detected on an annotated non-coding locus.

These results showcase the power of R2T2P in tracking gene expression products and their regulation from RNA to protein, while adding novel, massively upregulated species to the known DUX4-induced proteome.

### Beyond gene-level regulation from RNA to protein

While identifying novel transcribed gene loci compatible with protein synthesis, we also used the sub-gene routines of R2T2P to dissect DUX4-mediated isoform-level regulation in our dataset, again from RNA to Translation to Protein.

Using exon bin-level changes at 14 hours post-DUX4 induction, we detected thousands of genes with differential events in RNA-seq (**Figure 4A**, **Supplementary Table 5**), distributed along different genomic features. Although less numerous, Ribo-seq-derived events were more common in CDS regions, and RNA-seq-derived events also in 3’UTRs, highlighting the importance of tracking regulation across different steps in the RNA life cycle. Few significant events were specific to features in novel transcripts, with one example shown in **Figure 4B** in the *SELENOT* locus, again highlighting the importance of augmenting known gene models with custom annotation. When comparing the significance of sub-gene differential events against gene-level differential analysis (**Methods**), we observed a clear separation (**Figure 4C**), suggesting potentially distinct molecular mechanisms underlying multimodal DUX4-mediated regulation. This was visible using either Ribo-seq or RNA-seq significance values (**Supplementary Figure 9**). Given the numerous sub-gene differential events in the Ribo-seq, we queried our mass spectrometry dataset to test for the presence and regulation of alternative protein isoforms. We calculated a *deltaFC* coefficient, measuring the difference, for each peptide, between its up-or down-regulation compared to other peptides from the same gene, a strategy akin to *DEXseq* or other exon-level differential analysis strategies for sequencing data (**Methods**). Comparing *deltaFC* between peptides coming from up-or down-regulated exons, we observed a clear concordance (**Figure 4D**), with peptides from down-regulated exons being less abundant than other peptides from the same gene, or more abundant when coming from up-regulated exons. When using RNA-seq instead of Ribo-seq to define regulated exons, this trend was present albeit less pronounced, again confirming the importance of using Ribo-seq to quantify differences in active translation events. To account for differences in sensitivity between Ribo-seq and RNA-seq-based exon-level differential analysis (**Figure 4A**), we used the top 200 differential events, which showed a stronger agreement between upregulated sub-gene regions and their corresponding *deltaFC* values (**Supplementary Figure 10**). One example of coordinated sub-gene level regulation is shown in Figure 4E, where a differential event is detected in the *RTN4* locus, resulting in the expression of an alternative transcript, translated ORF, and protein isoform following *DUX4* activation.

**Figure 4:**
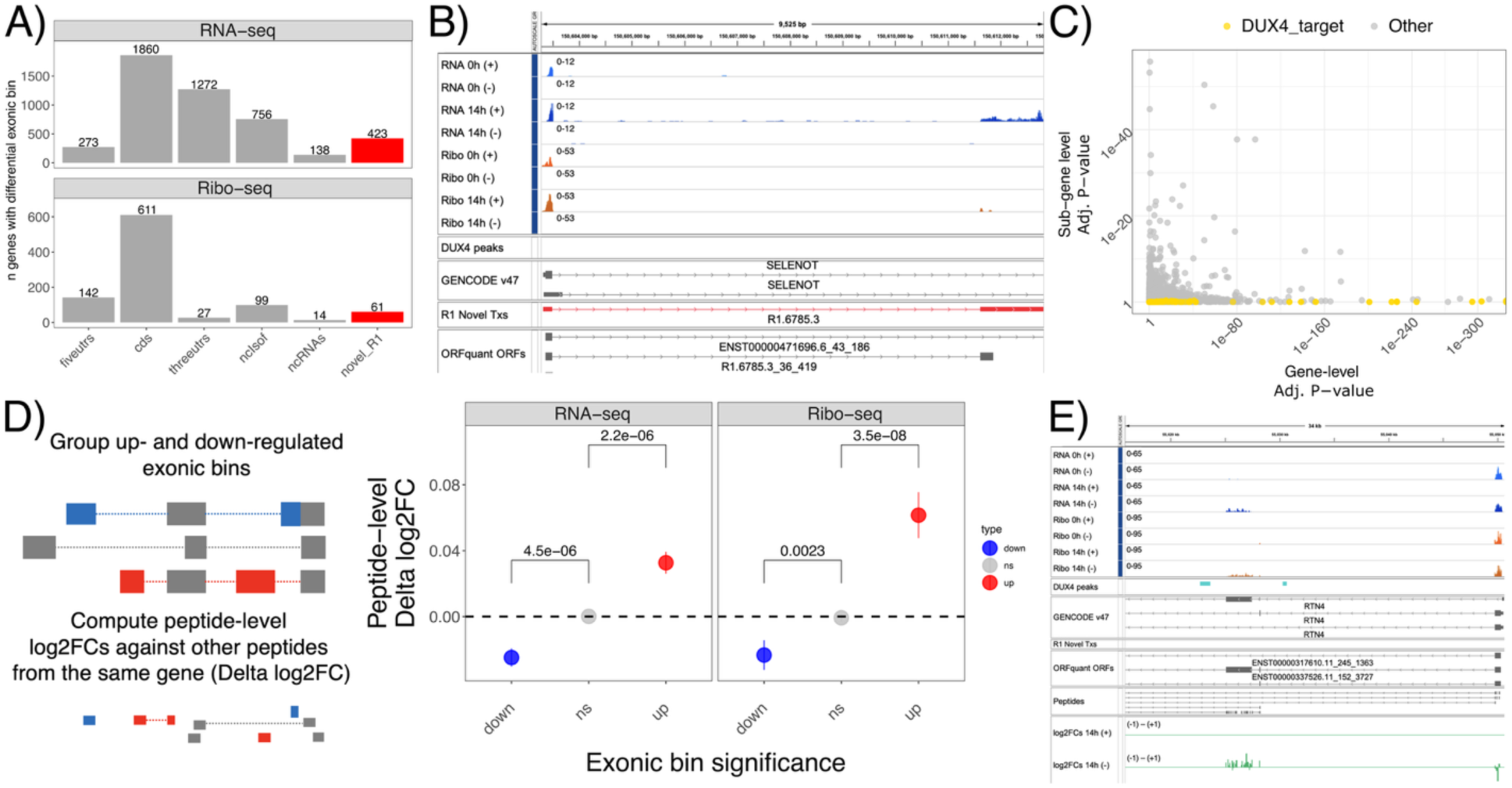
Differential analysis at the sub-gene level from RNA to Protein. In A) number of genes harboring differential events (from *DEXseq*) across multiple transcript regions, using RNA-seq or Ribo-seq at 14h post-DUX4 induction. In B) an upregulated novel exon in the *SELENOT* locus. In C) P-values from gene-level analysis using *DESeq2* (x-axis) against P-values derived from transcript features (exonic bins and junctions) from *DEXseq* (y-axis), using Ribo-seq. In D) a schematic depicting our analysis strategy, with delta log2FC of peptide abundances (using log2FC from other peptides of the same genes) divided according to the regulation of their originating exonic bin, using RNA-seq or Ribo-seq-derived significant transcript features (adjusted p-value < .05). P-values from paired Wilcoxon test against values from peptides coming from non-significant exons are shown. In E) an example over the *RTN4* locus, showing an alternative transcript being upregulated in RNA-seq, Ribo-seq and proteomics.

Taken together, these results show the power of *R2T2P* in detecting expression changes well beyond gene-level summarizations, and in uncovering robust regulation of the alternative transcriptome and its functional consequences to translation and the cellular proteome.

### An augmented long-read derived transcriptome further enhances identification and quantification

Given the increasing availability of custom transcriptome annotations for a variety of samples, we included in R2T2P the possibility of using an additional transcriptome annotation (**Figure 1**), increasing the complexity of the set of transcript structures considered throughout the entire workflow (**Discussion**). We decided to monitor changes in identification of differential events in our dataset using a recent Iso-seq-derived annotation in a similar DUX4-induction system in human muscle cells^36^.

The long-read-derived annotation comprises several novel genes and isoforms, with similar categories identified by our RNA-seq derived assembly (**Supplementary Figure 11**). When merging all the annotations (**Methods**), we derived a set of genes (**Figure 5A**) unique to the short-read-derived annotation (R1), unique to the long-read-derived annotation (R2), or shared between them (R1R2).

**Figure 5:**
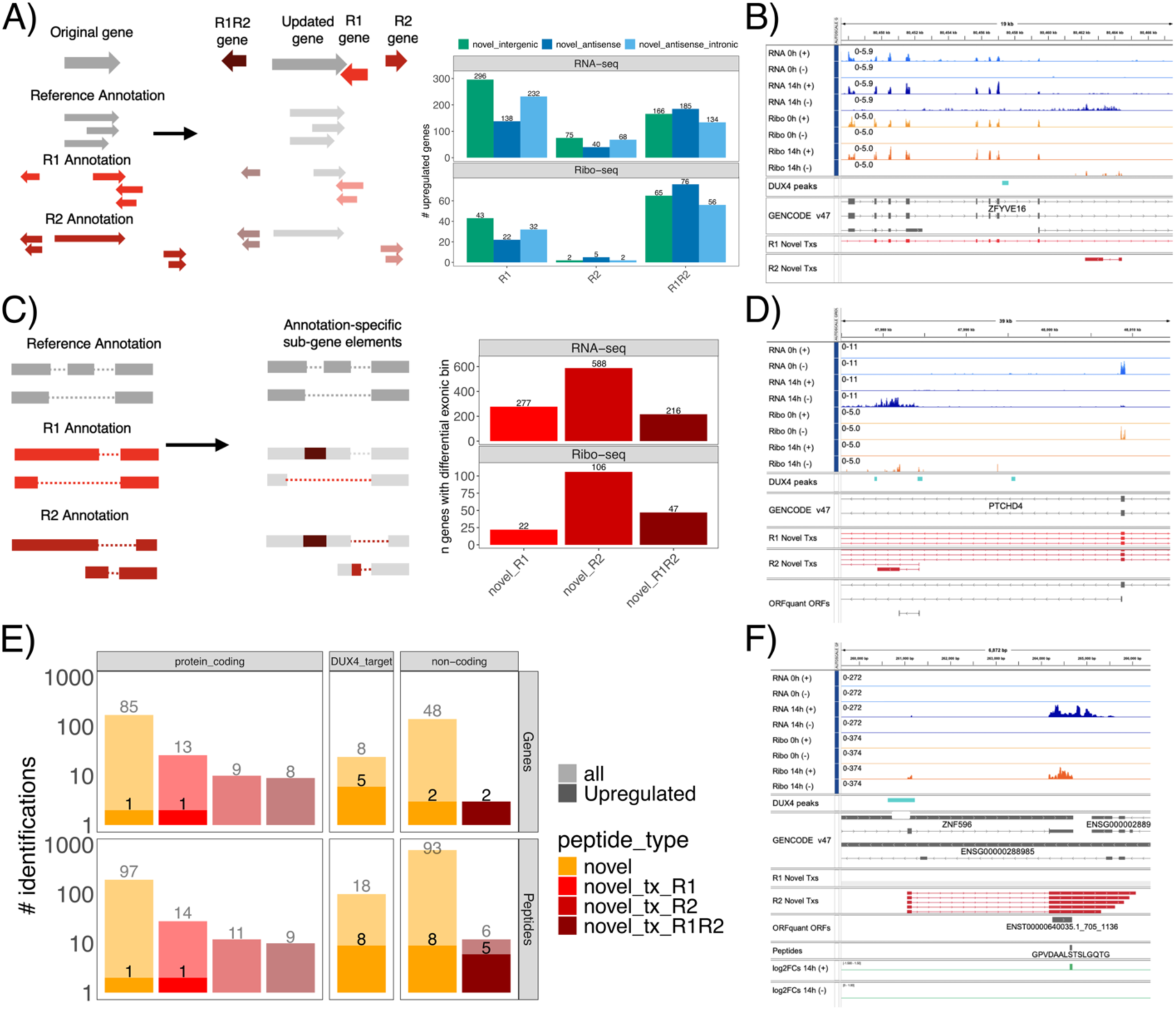
Assessing the contribution of multiple annotation sources with R2T2P. In A) a scheme on updating gene models using multiple annotations, with resulting genes unique to one annotation or comprising transcripts from multiple sources (left). On the right, results from gene-level differential analyses, using RNA-seq or Ribo-seq, with an example antisense intronic RNA shown in B). In C) a cartoon showcasing the strategy to obtain annotation-specific sub-gene features, alongside results from differential sub-gene analysis from *DEXSeq* at 14h; an example isoform is shown in D). In E) results at the level of peptides and genes from mass spectrometry, highlighting the number of differential species detected from different annotations. In F) the *ZNF596* locus, showing novel transcripts identified only by R2 annotation, and able to recapitulate coverage changes in RNA-seq and Ribo-seq, as well as a novel upregulated peptide.

At gene level, most differential events in our datasets belonged to shared or short-read-only derived annotation, especially at the level of translation (**Figure 5A**), with R2-only and R1-only genes being mostly upregulated at the level of RNA. One example of R2-only regulated gene is shown in **Figure 5B**. At the sub-gene level (**Figure 5C**), the R2 annotation allowed to detect hundreds of additional genes with sub-gene DE events in both RNA-seq and Ribo-seq (∼600 and ∼100, respectively), showing the power of long reads in pinpointing novel isoform-level changes, albeit often for lowly expressed RNAs (**Figure 5D**). With respect to ORF identification, we detected thousands of additional translated ORFs unique to the R2 annotation, albeit mostly as lowly translated novel isoforms of coding genes (**Supplementary Figure 12**). With respect to the Protein module, peptide-level differential analysis mostly supported peptides identified by both annotations, with few identifications unique to either annotation (**Figure 5E**) and almost none significantly upregulated. However, while our stringent analysis did not show upregulated peptides uniquely mapping to R2 transcripts only (**Methods**), this additional transcriptome allows to more precisely monitor gene expression and regulation. As an example (**Figure 5F**), in the *ZNF596* locus we identified a novel peptide mapping to a 3’UTR of a transcript; while initially identifiable as translation in a 3’UTR, R2 annotation (further supported by DUX4 binding evidence) shows a novel internal transcription start site, which can better explain the observed changes in RNA-seq, Ribo-seq, and peptide-level signals, thus offering an alternative, more straightforward explanation without the need to consider the complex dynamics of ribosome recycling and reinitiation (**Discussion**).

In summary, by including this additional step in the R2T2P workflow, we could improve our ability to detect otherwise invisible changes in our data, showcasing the importance of employing precise, custom transcriptome annotations.

### A novel DUX4-induced transcriptome in FSHD patients

Using our custom transcriptome annotation in iDUX4 cells, we wanted to assess its ability to identify molecular changes closer to the FSHD disease state. To this end, we used R2T2P to re-analyze RNA-seq data coming from patient-derived myoblasts differentiated into myotubes^32^, as well as RNA-seq data derived from muscle biopsies of patients stratified by different DUX4 markers’ expression and correlated with disease severity^33^ (**Figure 6A**). We detected significant upregulation of thousands of genes (**Supplementary Figure 13**), including a number of novel genes comparable to known DUX4 targets (**Figure 6A**, **Supplementary Table 6**) for both patient-derived myotubes and patient biopsies, effectively doubling the number of putative DUX4 target genes detected in FSHD patients. Among the novel upregulated loci (**Figure 6B**), we identified the same 2 novel protein-coding loci shown in **Figure 3D**. At the sub-gene level, we could again detect significant changes in hundreds of genes, often not correlated to gene-level changes (**Figure 6C, Supplementary Table 7**). Nonetheless, we detected a substantial overlap in sub-gene-level regulated genes between the 3 datasets (**Figure 6D**), including a common regulated sub-gene event in the *ZNF596* locus (**Figure 6E**), the same locus shown in **Figure 5F**.

**Figure 6:**
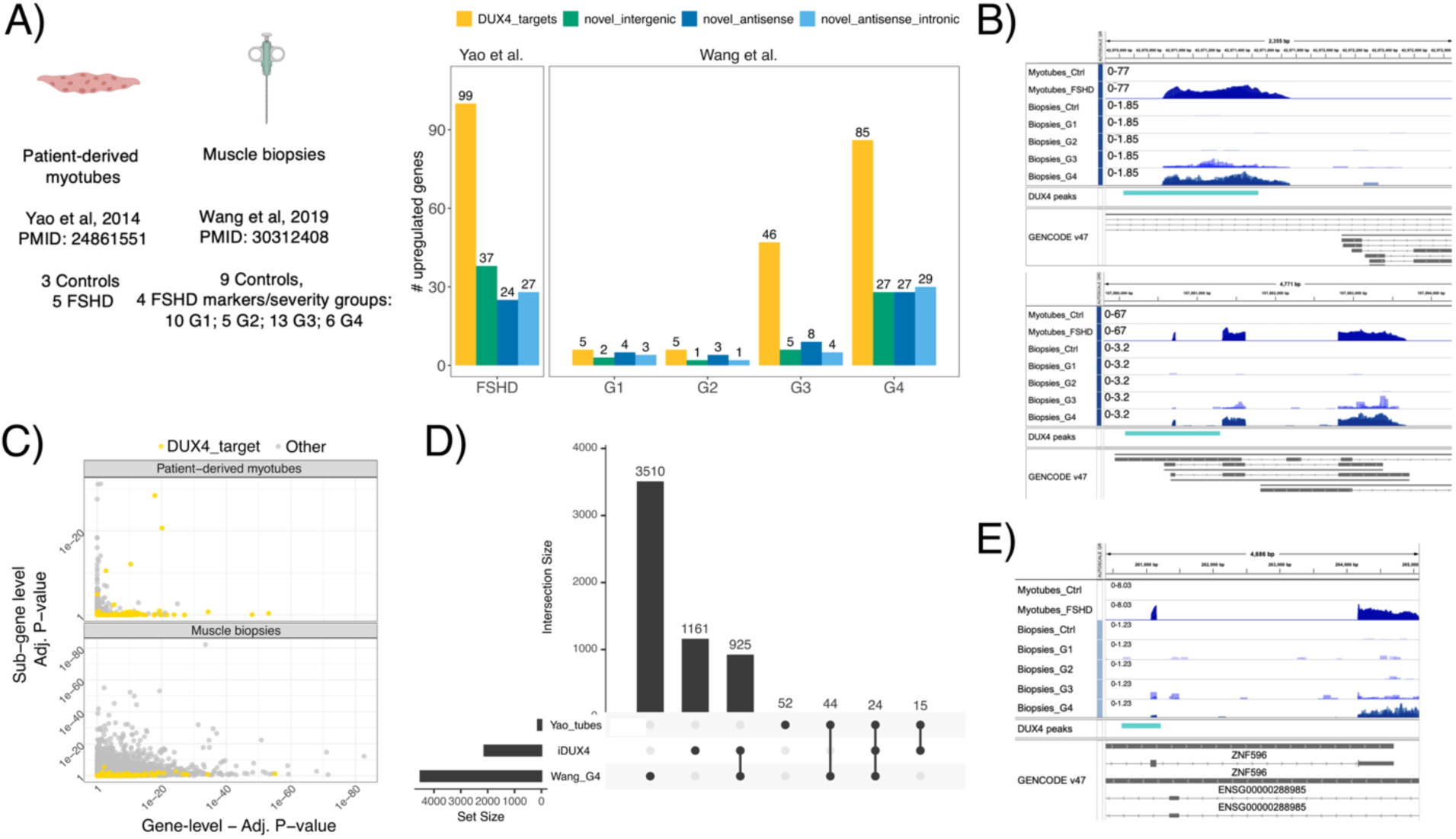
R2T2P-derived annotation reveals new targets in FSHD patients. In A) schematics of the datasets used (left), with gene-level differential analysis results (right). In B) the same examples as in Figure 3, showing expression directly from the patient datasets. In C) sub-gene-level analyses as in Figure 4, using RNA-seq from the two datasets. In D) an Upset plot showing the overlap between sub-gene-level regulated genes in iDUX4 cells, patient-derived myotubes and muscle biopsies, with an example sub-gene differential event in the *ZNF956* locus shown in E).

In summary, we confirmed that R2T2P-identified changes identified in iDUX4 cells are disease relevant, and expand the known molecular landscape of FSHD by further incorporating an alternative isoform dimension into the diseased cellular state, ultimately broadening the therapeutic landscape for this rare disease.

## Discussion

In this work, we presented R2T2P, our computational platform combining genome-guided transcriptome discovery and isoform-level *de novo* ORF finding with proteome-scale identification and quantification. Beyond providing a unique integrative system that tracks expression and regulation across multiple steps, R2T2P significantly improves traditional data analysis approaches that rely only on reference annotations and reference protein databases: it builds custom transcriptomes and translational landscapes, as well as custom protein databases. In its current implementation, our approach reduces reliance on protein inference, and uses peptide-level information to quantify changes in the proteome, unlocking the ability to perform protein isoform-aware differential analysis. Importantly, R2T2P allows for specific tracking and differential expression of novel species; by shifting the focus from validating their existence to quantifying their regulation across experimental conditions, we believe our approach represents an important advancement in discovery-oriented research. We envision such novelty-aware approaches to become routine in modern -omics based projects, where researchers can fully exploit the high quality and resolution of their data to build personalized reference transcriptomes, as well as translatomes and proteomes. Such a strategy holds great value for a wide range of biological applications, from annotation of non-model organisms to personalized medicine.

Remarkably, we effectively doubled the number of DUX4 target genes and validated the protein-coding capability for dozens of them. This was achieved despite the inherent sensitivity limitations associated with profiling the steady-state proteome, as opposed to employing potentially more sensitive approaches designed to capture rapid expression dynamics in the nascent proteome^57^. First identified in our iDUX4 dataset, we successfully confirmed the presence of novel RNAs in patients’ muscle biopsies, despite the challenges posed by their complex cellular composition. This validation carries significant translational weight, as it expands the repertoire of potential mechanistic disease biomarkers and therapeutic targets, a critical asset for rare diseases where options are frequently limited.

However, increasing the breadth of the reference annotation inherently introduces ambiguity when treating “genes” as rigid entities whose function is to encode a single protein that dictates them their name or symbol. Given the presence of diverse, and regulated, RNA and protein products arising from a single genomic locus, we encourage researchers to fully embrace the functional heterogeneity of gene expression output rather than rely on oversimplified, gene-centric frameworks that favour reductive interpretations.

Moreover, while we provide gene-level results, the frequent occurrence of overlapping transcribed loci (**Figure 2**) complicates interpretation of the precise function of gene expression products. This becomes even more intricate when using multiple annotations. Pseudoalignment-based quantification tools for RNA-seq can provide transcript-level quantification estimates to resolve such complex scenarios^58,59^, and potentially allow to discard many lowly expressed RNAs. Inclusion of such approaches, albeit potentially sensitive to different parameters^60^, might represent a future direction to filter transcript annotations.

In our hands, including an external custom annotation improved the identification of novel species, while not adding a substantial number of strongly upregulated RNAs and proteins. However, long-read technologies provide a more faithful representation of the transcriptome, improving the interpretation at the isoform-level, in our case pointing to regulated internal transcription start sites in both iDUX4 cells and FSHD patients (**Figure 5F** and **Figure 6E**).

DUX4 is known to induce transcription at repetitive elements during early embryonic development, with a substantial overlap between target regions in development and in FSHD muscle^61^. The precise identification and quantification of these elements remains challenging, especially given the short nature of RNA-seq and Ribo-seq data, and peptides from bottom-up mass spectrometry methods. While showing the advantages of using an additional long-reads derived annotation, the findings reported in this study all come from using uniquely mapping reads, likely reflecting only a subset of DUX4-induced changes. Importantly, R2T2P allows for the use of all or uniquely mapped reads for differential analyses or ORF finding, allowing researchers to compare results using different parameters.

The core strength of the R2T2P workflow lies in its ability to track heterogeneity in expression at sub-gene level. This dimension is often overlooked in many single-cell and spatial -omics approaches, where, driven by resolution constraints and to streamline analysis and interpretation, popular pipelines often rely on a “*one-gene, one-protein”* assumption. Importantly, we do not assume a unique mechanism underlying sub-gene level changes: the presence of DUX4 binding sites led us to hypothesize alternative transcription initiation as a source of many such events^62^ ; changes in 3’UTRs (**Figure 4A**) might suggest alternative polyadenylation, while regulation of non-coding isoforms is indicative of nonsense-mediated decay regulation, which we quantified and examined in a previous study^34^. Despite the vast complexity of the transcriptome, our high-quality Ribo-seq allowed us to focus on cytoplasmic RNAs, effectively filtering out most nuclear-retained species and quantifying diversification of the synthesized proteome versus other aspects of gene expression regulation.

Implemented in Nextflow to maximize reproducibility and cross-platform interoperability, we envision leveraging the R2T2P framework as the foundational architecture for future modular expansion. These will include dedicated sub-modules to quantify mRNA and translation regulation, integrated support for additional proteomics workflows via *FragPipe*, and add compatibility with *de novo* peptide sequencing algorithms^63,64^, towards a fully unbiased platform for RNA and protein integration.

## Materials and Methods

### Initial RNA-seq mapping and junction discovery

*STAR 2.7.9a* is used to align RNA-seq reads to the genome. Two different types of RNA-seq alignments are performed.

In the first type of alignments, which are hereafter referred to as two-pass alignments, STAR is run in two-pass mode (*--twopassMode Basic*) and the Gene Transfer Format (GTF) file of a reference transcriptome annotation is provided to the STAR parameter *--sjdbGTFfile*. Standard local alignment with soft-clipping allowed is used (*--alignEndsType Local*). Alignments are performed allowing up to 4 mismatches *(--outFilterMismatchNmax 4*) and allowing up to 250 alignments for a read *(--outFilterMultimapNmax 250*). Regarding chimeric alignments, values for different STAR parameters were specified to have a minimum segment length of 15 (*--chimSegmentMin 15*), a minimum junction overhang of 10 (*--chimJunctionOverhangMin 10*), a minimum total score of chimeric segments of 20 (*--chimScoreMin 20*), and a minimum difference between the best chimeric score and the next one of 10 (*--chimScoreSeparation 10*). The Sequence Alignment Map (SAM) attributes NH, HI, AS, nM, NM, MD, and XS are included in the sorted BAM output (*--outSAMattributes NH HI AS nM NM MD XS*; *--outSAMtype BAMSortedByCoordinate*). Separate output files contain unmapped and partially mapped reads, i.e., mapped only one mate of a paired end read (*--outReadsUnmapped Fastx*). No mismatches for stitching of the splice junctions were allowed for non-canonical motifs, GT/AG and CT/AC motif, GC/AG and CT/GC motif, and AT/AC and GT/AT motif (*--alignSJstitchMismatchNmax 0 0 0 0*). A single output alignment is reported for each multimapping read (*--outSAMmultNMax 1*), and the *Old_2.4* option is specified for the ordering of alignments of multimapping reads (*--outMultimapperOrder Old_2.4*). The random number generator seed is set to 0 (*--runRNGseed 0*). The intron motif is used to derive the strand (*--outSAMstrandField intronMotif*). Multithreading is used, as STAR is run with 12 threads (*--runThreadN 12*).

In the second RNA-seq mapping, a reference transcriptome annotation, as well as novel junctions discovered in the first mapping, are provided to STAR, using the parameters *-- sjdbGTFfile* and *--sjdbFileChrStartEnd*, respectively. A minimum overhang of 500 nt is specified for unannotated junctions (*--alignSJoverhangMin 500*) to disable novel junction discovery, and up to 5,000,000 junctions can be inserted on the fly *(--limitSjdbInsertNsj 5000000*). Other parameters were kept identical to the first mapping step.

### Transcriptome reconstruction

In the RNA module the assembler *StringTie3* is used for reference-guided transcriptome reconstruction, using GENCODE v47 as the reference transcriptome. The assembler is run by using the conservative mode and by setting the minimum length for the predicted transcripts to 100. RNA-seq alignments from the second mapping step and a reference transcriptome annotation file are provided to *StringTie3*.

### Combining transcriptome annotations

A merging function takes as input the reference transcriptome annotation and a *GFFCompare* output GTF file, which is generated by comparing the *StringTie3* GTF file (R1) to the reference annotation. *GFFCompare* class codes are used to classify *StringTie3*-identified transcripts according to their positions with respect to reference transcripts (e.g., identical, contained, intronic, antisense, and intergenic). Gene and transcript biotypes are also assigned accordingly. *StringTie3*-assembled transcripts which are present in the reference annotation (i.e., *StringTie3* transcripts with *GFFCompare* class codes “=” and “c”) are not included in the result of the merging. The script produces a single annotation file which contains reference transcripts as well as novel *StringTie3*-assembled transcripts.

When using an additional annotation, an extra step takes as input the *StringTie3* transcriptome annotation and a *GFFCompare* output GTF file, which is generated by comparing the user-provided annotation (R2) to the *StringTie3* (R1) annotation. Notably, user-provided transcripts which are present in the *StringTie3* annotation (i.e., user-provided transcripts with *GFFCompare* class codes “=” and “c”) are here included in the final merged set, to avoid assignment of similar transcripts to one reference only. The script produces a GTF file which contains coordinates of *StringTie3* (R1) and user-provided (R2) transcripts. This merged R1R2 annotation is then compared to the reference annotation again with *GFFcompare*, as described in the first step.

### Ribo-seq pre-processing

Adapter sequences were removed with *cutadapt*, identical Ribo-seq reads were collapsed to FASTA format with *fastx_collapser*, and Unique Molecular Identifiers (UMIs) were also removed. After removing contaminant sequences (rRNAs, snoRNAs and miRNAs, tRNAs) through a series of alignments with *Bowtie2*, the remaining Ribo-seq sequences were stored in FASTA files.

### Final mapping of RNA-seq and Ribo-seq reads

The output GTF file of the RNA module is provided by using the STAR parameter *-- sjdbGTFfile*. Novel junction discovery is disabled by specifying 500 as the minimum overhang for unannotated junctions (*--alignSJoverhangMin 500*). For the local alignment, SAM attributes, and strand information from the intron motif, the same parameter values as in the two-pass RNA-seq alignments are specified. Parameters for chimeric alignments have the same values as in previous alignments, except for parameter *-- chimJunctionOverhangMin*, which is kept at its default value. In these final alignments, some STAR parameters have different values depending on the specific assay type of the provided data (RNA-seq or Ribo-seq): up to 4 mismatches per alignment are allowed for RNA-seq data, but up to 3 for Ribo-seq data (parameter *--outFilterMismatchNmax*); the maximum number of allowed alignments for a read is 250 when using RNA-seq reads, whereas it is 50 when aligning Ribo-seq reads (parameter *--outFilterMultimapNmax*). Finally, as in previous alignment steps, there is only one reported alignment per multimapping read (*--outSAMmultNmax 1*). The Bioconductor R packages *GenomicRanges*, *GenomicFeatures*, *GenomicAlignments*, *rtracklayer* and *RiboseQC* were used for downstream processing of the alignment files.

### Gene-level and sub-gene level differential analyses

Read counts on exons (for all genes) and CDSs (for protein-coding genes) were used to perform gene-level differential analyses with the R package *DESeq2*, whereas counts on flattened gene models, consisting of individual exonic bins, junctions, and introns for each gene, were used to perform sub-gene differential analyses with *DEXSeq*. For each differential analysis, uniquely mapping reads were used. Upregulated genes were selected using a log2FC>0 and an adjusted p-value threshold of 0.05. Sub-gene significant events were selected using an adjusted p-value threshold of 0.05. To obtain values shown in Figure 4C and Figure 6C, P-values across sub-gene events are aggregated per gene by taking the minimum P-value from both exon bins and junction-level features.

### Filtering overlapping genes

To limit potential wrong identification of regulated genes due to overlapping genes, we applied the following strategy: genes with >50% of their RNA-seq reads mapping to a different gene were initially selected; for each of those genes, if no feature (a junction or an exon bin) unique to them was significantly regulated (as defined by *DESeq2*) those genes were removed. This added an additional column to all output files (*gene_overlapping_noDE*) which was used as a filter for all gene- and sub-gene level analysis shown in the paper.

### Additional differential analyses and post-processing

Changes at the level of translation are calculated using *DESeq2* by using assay type (RNA-seq or Ribo-seq) as an additional covariate in the model^65,66^; the same is performed using intronic or exonic counts to infer post-transcriptional regulation^67^ ; GO enrichment analysis is performed using results from each differential analysis (gene-level RNA, gene-level translation, sub-gene level, etc …), as well as training of regression models to predict changes in RNA or translation from different sequence features^68^. This comprehensive differential analysis strategy helps distinguishing changes in transcription, translation, as well as sub-gene regulatory events (such regulation of splicing, polyadenylation, or transcription initiation), while tracking signal from novel transcripts.

### Assessment of DUX4 binding with ChIP-seq data

Promoters were defined as genomic regions comprising 1000 nucleotides upstream of Transcription Start Sites (TSSs) and 300 nucleotides downstream. Genes were defined as supported by chromatin immunoprecipitation sequencing (ChIP-seq data) if at least one of their transcript promoters overlapped a peak. When focusing on promoters of a specific gene group, peaks overlapping promoters of the other gene groups were not considered for the analysis. The proportion of ChIP-seq-supported genes for each group was then calculated by dividing the number of supported genes by the total number of genes in the group.

### *De novo* detection of translated ORFs

The output transcriptome annotation of the RNA module as well as the Ribo-seq alignment results of the Translation module are provided to the R package *RiboseQC* to get Ribo-seq read counts. Then, the R package *ORFquant* is provided with the transcriptome annotation file and with a *RiboseQC* output file, using default parameters. By using *ORFquant* translated regions on transcripts are detected. Only the signal from uniquely mapping reads was used to detect translated ORFs on GENCODE v47, *StringTie3*, and long-read-derived transcripts.

### Database searches and peptide quantification

Protein sequences corresponding to CDSs of the reference transcriptome annotation are extracted with a custom R script. The script uses functions from the packages *GenomicFeatures*, *Biostrings*, and *GenomicRanges*. Annotated protein sequences are then saved to a FASTA file (*ref_database*). The same R script also saves a FASTA file containing protein sequences corresponding to *ORFquant*-detected ORFs (*custom_database*) as well as a FASTA file containing both sets of protein sequences (*combined_database*). Decoys and contaminants are then added to the three protein databases using *Philosopher v5.1.3-RC9*, which is included in the computational platform *FragPipe v24.0*. Then, database searches and peptide quantification are performed with *FragPipe v24.0*.

For the analysis of the iDUX4 data, the TMT dataset and the three protein databases were provided to *FragPipe v24.0*. The software tools *MSFragger v4.4.1* and *IonQuant v1.11.20* were used. Proteomic searches were run with the following settings: the basic TMT 16-plex workflow was used, with quantification and identification from MS2; Met oxidation, protein N-term Acetyl, and n-term TMT were specified as variable modifications; *Philosopher* filtering was used with its default value; *TMT-Integrator* was run by using the reference approach, doing data summarization at all levels, with no normalization or log2 transformation. When running the database search tool *MSFragger*, *stricttrypsin* was used as search enzyme, a minimum peptide length of 7 and a maximum length of 50 were specified, and up to 2 missed cleavages were allowed.

### TMT labeling mass spectrometry

Proliferating MB135 iDUX4 human myoblasts (PMID: 28171552) were cultured in Ham’s F-10 Nutrient Mix (Gibco) supplemented with 20% fetal bovine serum (Gibco), 10 ng/ml recombinant human basic fibroblast growth factor (Promega), 1 uM dexamethasone (Sigma), and 2 ug/ml puromycin (VWR). To induce DUX4 expression, 1 ug/ml of doxycycline (Sigma) was added for 4, 8, or 14 hours. Cells were lysed in RIPA Buffer (Thermo Scientific) containing protease inhibitors (Sigma) and subjected to proteomics analysis at the Fred Hutchinson Cancer Center Proteomics & Metabolomics Shared Resource.

A reference bridge sample was created by mixing 8.3 µg of protein from each sample into a single tube (300 µg total protein) and dividing it into three 100 µg aliquots. Protein lysates (36 total) from each sample consisting of 100 µg of protein along with the 3 reference bridge samples were reduced with 5 mM tris (2-carboxyethyl) phosphine (TCEP) at room temperature for 15 min and alkylated with 10 mM 2-chloroacetamide at room temperature for 30 min. A chloroform-methanol precipitation was carried out and protein pellets were resuspended in 70 µL 50 mM HEPES pH 8.5, digested with 1 µg Lys-C (Promega) at room temperature for 4 hours, and digested with 1 µg trypsin (Promega) at 37 °C overnight with shaking. TMTpro reagents (ThermoFisher Scientific) were resuspended in 30 µL of acetonitrile and added to the assigned sample or reference bridge. After labeling for 1 hour, the reference bridge aliquots were combined, mixed to heterogeneity, and divided into 70 µL aliquots. A reference bridge aliquot was assigned to each batch of TMT-labeled samples to form three total batches, each containing 12 TMT-labeled samples and 1 TMT-labeled reference bridge. A 2 uL aliquot from each member of the first batch was mixed 1:1 across all TMT channels, subjected to vacuum centrifugation to remove acetonitrile, desalted with a C18 Ultra-micro Spin Column (Harvard Apparatus), and analyzed as a “label check” with an Orbitrap Eclipse mass spectrometer. Based on the summed protein intensity for each TMT-channel in the label check, a volume from each member of the batch was mixed to equalize the amount of protein from each member contributing to the batch. The label check and mixing were repeated for the two additional batches. The batches were desalted on 3 cc/200 mg C18 SepPak cartridges (Waters) and dried by vacuum centrifugation.

### Basic reverse-phase (bRP) fractionation

The desalted TMT batches were resuspended in 100 µL of 10 mM ammonium bicarbonate pH 8. The material was loaded on to a Zorbax 2.1 cm x 150 mm (5-µm particle size) Extend-C18 column (Agilent) connected to a ThermoFisher Scientific Vanquish HPLC equipped with a diode array detector and fraction collector. The sample was gradient-eluted from the column at a flowrate of 250 µL/min over 55 minutes using a combination of solvents “A” (10 mM ammonium bicarbonate) and “B” (acetonitrile). The elution profile used was as follows: from 0 to 5 minutes “B” was held at 1%, from 5 to 55 minutes “B” varied from 5% to 40% followed by an increase to 90% “B” over 5 minutes, and then a hold for an additional 5 minutes at 90% “B”. The UV signal was monitored at 210 nm and fractions were collected every 50 seconds, resulting in 96 fractions. The fractions were combined into 24 pools by concatenating every 24^th^ fraction into a pool (pool 1 = fractions 1, 25, 49, and 73; pool 2 = fractions 2, 26, 50, and 74; etc). The pools were taken to dryness by vacuum centrifugation and subsequently analyzed by LC/MS.

### LC/MS

The dried basic reverse phase fractions were brought up in 2% acetonitrile in 0.1% formic acid (20 μL) and analyzed (1.5 μL) by LC/ESI MS/MS with a ThermoFisher Scientific Easy1200 nLC coupled to a tribrid Orbitrap Eclipse with FAIMS pro mass spectrometer. In-line de-salting was accomplished using a reversed-phase trap column (100 μm × 20 mm) packed with Magic C_18_AQ (5-μm, 200Å resin; Michrom Bioresources) followed by peptide separations on a reversed-phase column (75 μm × 270 mm) packed with ReproSil-Pur C_18_AQ (3-μm, 120Å resin; Dr. Maisch) directly mounted on the electrospray ion source. Chromatographic separation was carried out adjusting the elution gradient from 4% to 9% B (80% acetonitrile with 20% water and 0.1% formic acid) in 2 minutes, 9% to 25% B in 80 min, 25% to 44% B in 60 min, 44% to 63% B in 8 min, holding at 63% for 1 min, and 63% to 90% in 1 min. A flow rate of 300 nL/minute was used for chromatographic separations. The temperature of the chromatographic column was maintained at 40 ℃ using a heating blanket. A spray voltage of 2300 V was applied to the electrospray tip in-line with a FAIMS pro source using varied compensation voltages of -40 V, -60 V, -80 V while the Orbitrap Eclipse instrument was operated in the data-dependent mode. MS survey scans were in the Orbitrap (Normalized AGC target value 300%, resolution 120,000, and max injection time 50 ms) using a 3 sec cycle time and MS/MS spectra acquisition were also detected in the Orbitrap (Normalized AGC target value of 250%, resolution 50,000 and max injection time 100 ms) using higher energy collision-induced dissociation (HCD) activation with a collision energy of 38%.

### Peptide-level and gene-level proteomic analyses

Differential analyses at the levels of peptides and genes are performed with the R package *limpa*. First, raw intensities are imputed using *limpa* with default parameters, and differential analysis is performed using raw or imputed intensities.

Intensities can then be grouped by either gene id, protein combination (computing all possible mapping database entry combination per each peptide), or gene:novelty combination, using R1 annotation, R2, or both (i.e. GeneX_annotated, GeneX_novel_R1, GeneY_annotated, …).

When aggregating peptides by gene (or other combination), an additional differential analysis is performed using the imputation fit by *limpa*, thus obtaining 3 different results for each comparison. Using peptide-level fold changes, *deltaFC* values are computed using the difference between each peptide fold change vs. the average of the other peptide fold changes, obtaining an estimate of peptide-level regulation different from other peptides from the same gene. All results shown in the paper come from using raw, not-imputed intensities.

### Identification of annotation-specific peptides

To rule out potential inconsistencies, each peptide is re-mapped back to annotated CDS and transcript regions. Peptides only assigned to entries from either R1 or R2 annotation, but mapping to annotated exons, are then considered “novel” (as they don’t map to entries in the reference database) but not unique to either annotation, one example being the one showed in Figure 5F.

## Supplementary Material

Supplementary Table 1. Summary statistics of the sequencing datasets used.

Supplementary Table 2. Gene-level differential analysis results in iDUX4 cells.

Supplementary Table 3. Transcript-level coordinates of ORFquant-detected ORFs in iDUX4 cells.

Supplementary Table 4. Detected peptides and genes in TMT data from iDUX4 cells.

Supplementary Table 5. Sub-gene differential analysis results in iDUX4 cells.

Supplementary Table 6. Gene-level differential analysis results in the analyzed FSHD datasets.

Supplementary Table 7. Sub-gene differential analysis results in the analyzed FSHD datasets.

Supplementary Figure 1 – Example execution times and required computational resources for each R2T2P step.

Supplementary Figure 2 – New identified transcripts across multiple biotypes.

Supplementary Figure 3 – Gene-level DUX4-mediated upregulation across different biotypes.

Supplementary Figure 4 – Robust upregulation of novel genes in both RNA-seq and Ribo-seq.

Supplementary Figure 5 – DUX4 binding enrichment at promoters of regulated genes.

Supplementary Figure 6 – ORFquant results on an augmented transcriptome.

Supplementary Figure 7 – ORFquant results using all or uniquely mapping Ribo-seq reads.

Supplementary Figure 8 – Peptide identification across protein databases.

Supplementary Figure 9 – Gene- and sub-gene-level regulation in Ribo-seq and RNA-seq.

Supplementary Figure 10 – Sub-gene agreement between RNA and protein using the top 200 DE bins.

Supplementary Figure 11 – Novel transcripts across annotations and gene biotypes.

Supplementary Figure 12 – Novel detected ORFs across annotations and gene biotypes.

Supplementary Figure 13 – Gene-level differential analysis in FSHD.

## Data availability

The RNA-seq and Ribo-seq dataset of iDUX4 cells are available at GEO (GEO Accession: GSE178761). DUX4 ChIP-seq peaks were taken from (L. N. Geng et al., *Developmental Cell*, 2012). Known DUX4 targets were taken from (Z. Yao et al., *Human Molecular Genetics*, 2014). The RNA-seq data of patient-derived and control myotubes is from (Z. Yao et al., *Human Molecular Genetics*, 2014), whereas the RNA-seq dataset of patient-derived biopsies and G1 controls is from (L. H. Wang et al., *Human Molecular Genetics*, 2019). The Iso-seq annotation (D. Zheng et al., *Sci. Adv.,* 2025) was kindly provided by the van der Maarel lab. RAW files for the TMT-labeling mass spectrometry dataset are available at MassIVE under accession MSV000103781.

Scripts to reproduce all figures are available at https://github.com/RobAlbn/R2T2P-Figures.

## Code availability

R2T2P is freely available at https://github.com/nfdata-omics/R2T2P/

## Author Contribution

R.A. built the entire R2T2P workflow, and performed all the analysis, supervised by L.C.. M.B. and C.J. supported the Nextflow implementation, with supervision from A.R. and L.C.. A.E.C. performed the TMT-labelling mass spectrometry experiment, supervised by S.J.. S.J. and G.T. provided important feedback and interpretation on DUX4- and FSHD-related findings. A.G. provided early support for proteomics data analysis, later continued by A.S.A. with supervision from J.K.. L.C. and R.A. wrote the manuscript, with important contributions from all Authors.

## Acknowledgments

The Authors want to thank the van der Maarel lab for kindly providing the Iso-seq-derived annotation for the DUX4-induced transcriptome. Additionally, L.C. and R.A. want to thank Dr. Fengchao Yu and Prof. Alexey I. Nesvizhskii for their kind FragPipe-related support. This work was supported by the University of Colorado School of Medicine Translational Research Scholars Program (S.J.), the NIH/NIAMS grant R01AR083381 (S.J. and A.E.C), and the NIH P30 CA015704 of the Fred Hutch/University of Washington/Seattle Children’s Cancer Consortium, which includes the Proteomics and Metabolomics Shared Resource, RRID:SCR_022618. GT is supported by the Academy of Medical Sciences Professorship Scheme (Grant APR8\1017).

