## Supplementary_Figures 1-13 for "Accelerating discovery and integration from RNA to Protein with *R2T2P*"

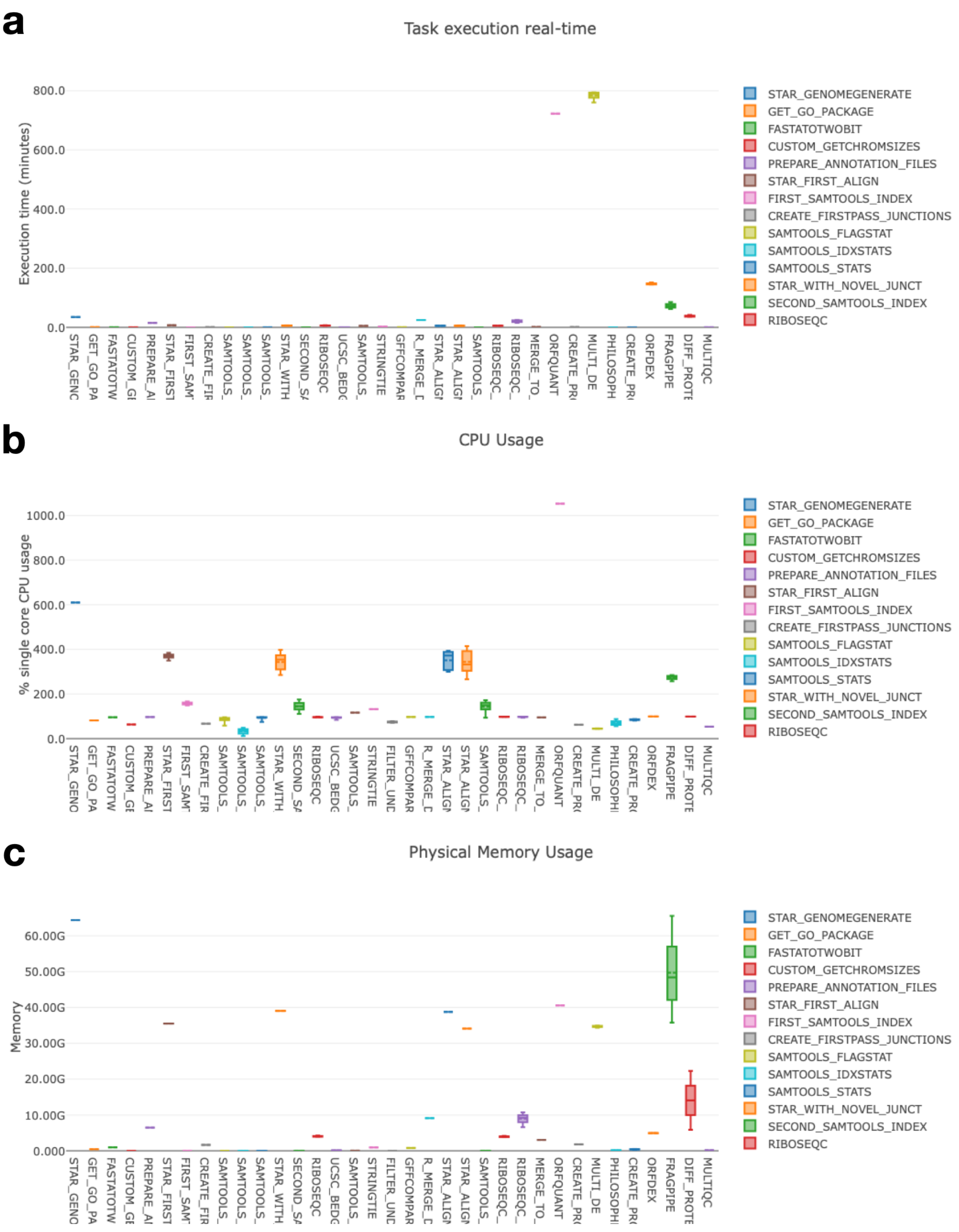


**Supplementary Figure 1** – Example execution times and required computational resources for each R2T2P step.

Execution times (a), percentage of single core CPU (Central Processing Unit) usage (b), and memory usage (c) for all the steps of the Nextflow pipeline, when applying it to an example dataset. The plots are included in the Nextflow execution report of the pipeline.

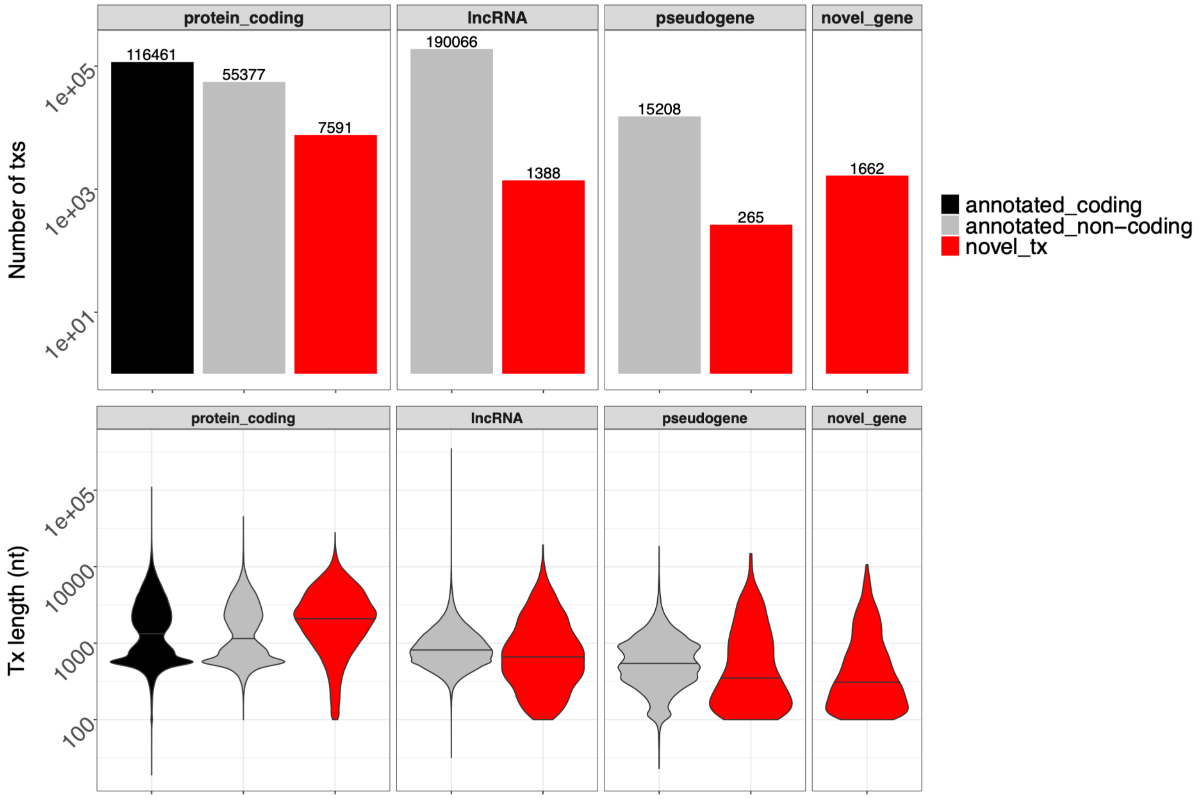


**Supplementary Figure 2** – New identified transcripts across multiple biotypes.

Number of annotated and novel StringTie-assembled transcripts are shown above, with their lengths in nucleotides shown below.

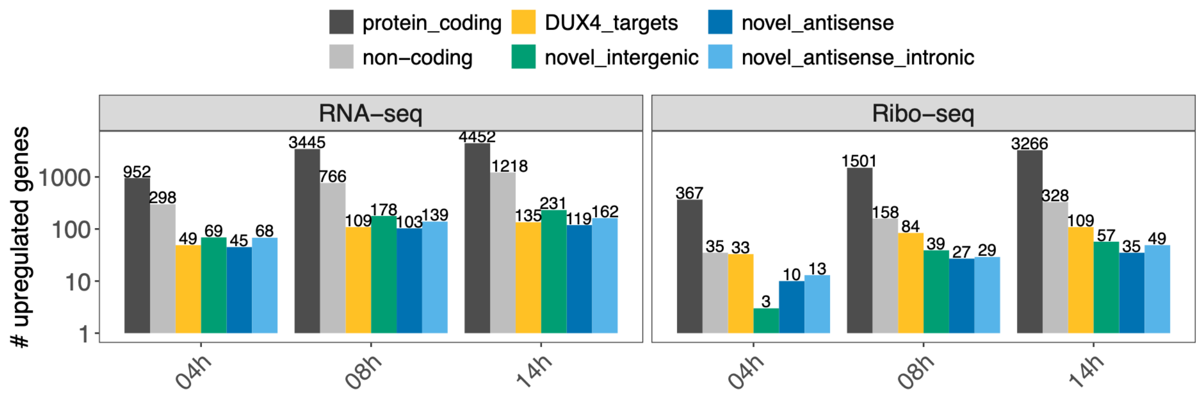


**Supplementary Figure 3** – Gene-level DUX4-mediated upregulation across different biotypes.

Number of upregulated genes across multiple annotation categories using either RNA-seq or Ribo-seq.

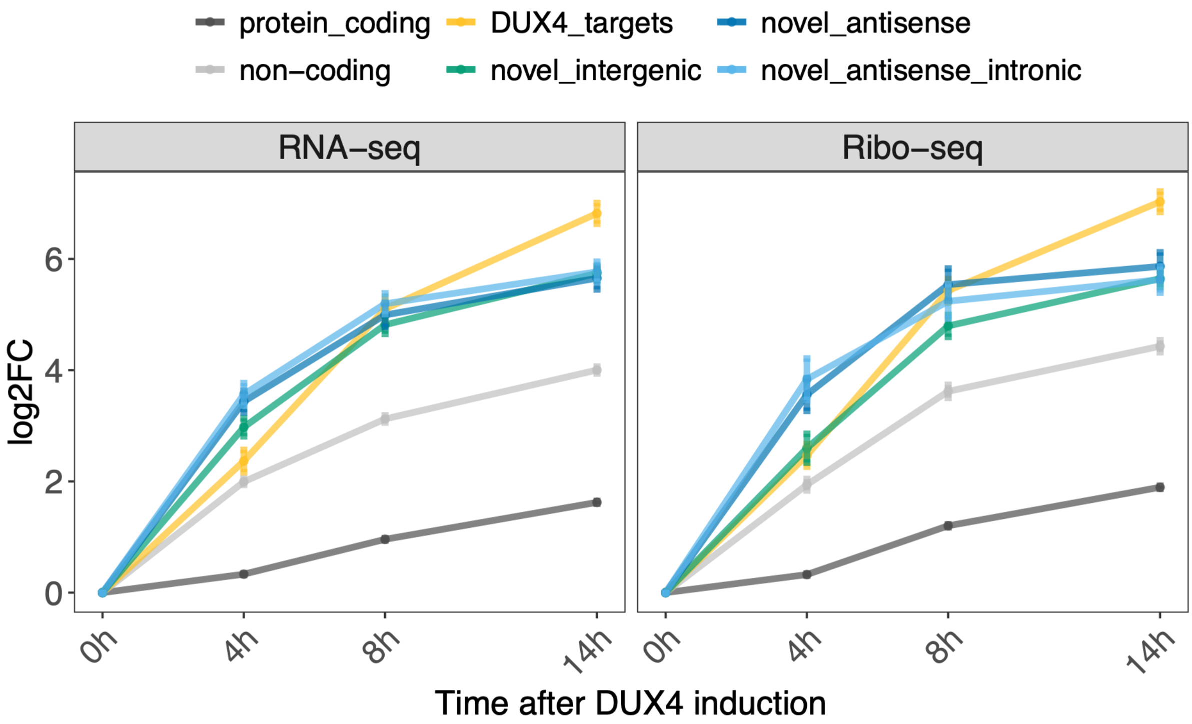


**Supplementary Figure 4** – Robust upregulation of novel genes in both RNA-seq and Ribo-seq.

Log2FC values for all upregulated gene categories, calculated using either RNA-seq or Ribo-seq data.

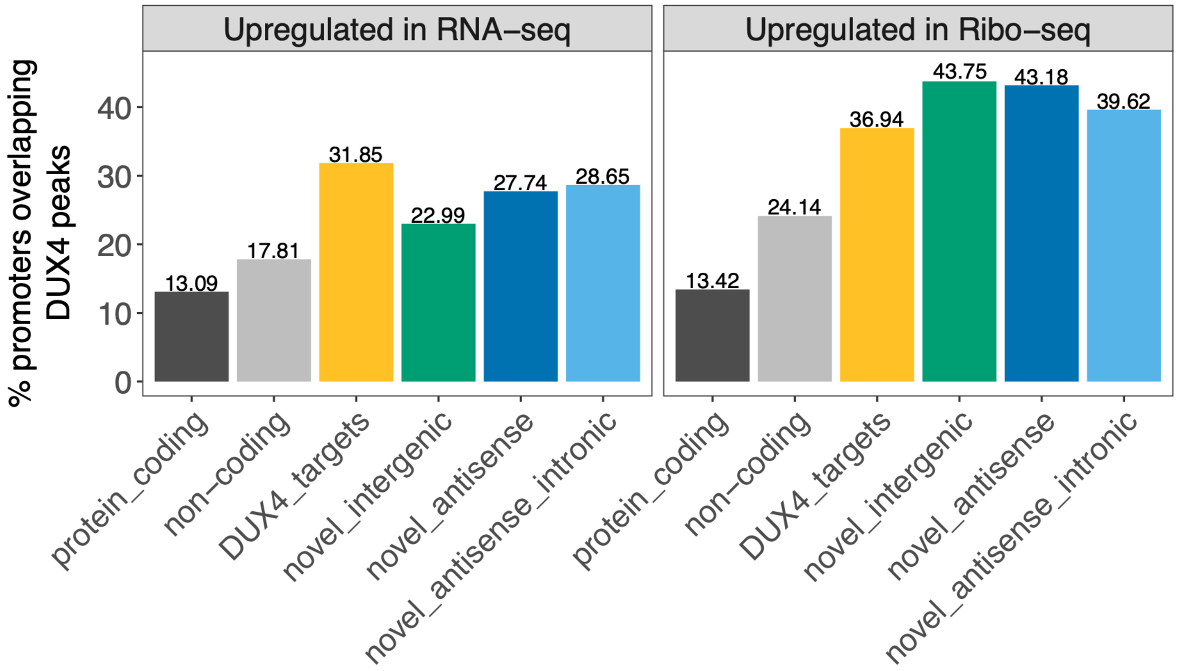


**Supplementary Figure 5** – DUX4 binding enrichment at promoters of regulated genes.

Percentage of promoters overlapping published DUX4 binding sites, using different gene categories upregulated at the level of RNA-seq or Ribo-seq.

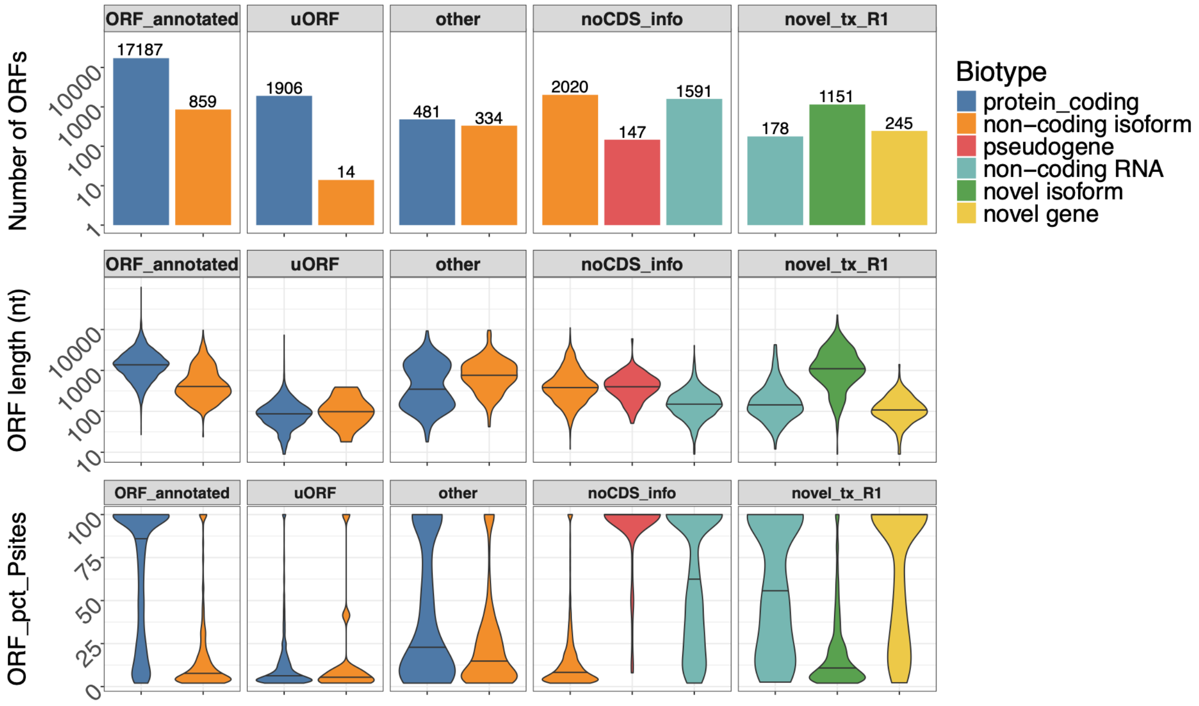


**Supplementary Figure 6** – ORFquant results on an augmented transcriptome.

Number of ORFs, ORF length in nucleotides, and percentage of gene translation are shown, stratified by ORF category and annotated biotype. “ORF_annotated” represents ORFs whose structure perfectly matches the annotated CDS; “other” represents additional ORFs, such as nested ORFs, overlapping ORFs, downstream ORFs in 3′ UTRs (dORFs), “noCDS_info” represents ORFs in transcripts with no CDS annotation, while “novel_tx_R1” represents ORFs uniquely mapping to novel transcripts. The maximum width of each violin plot is the same for each panel, and the median value is shown as a black bar.

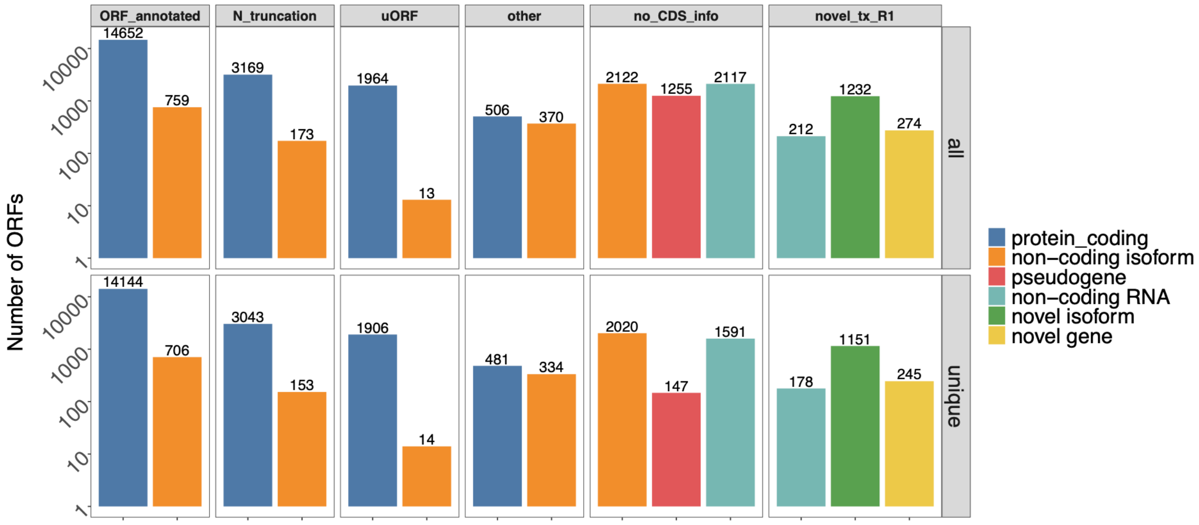


**Supplementary Figure 7** – ORFquant results using all or uniquely mapping Ribo-seq reads.

Number of detected ORFs across categories using only uniquely mapping reads or all reads.

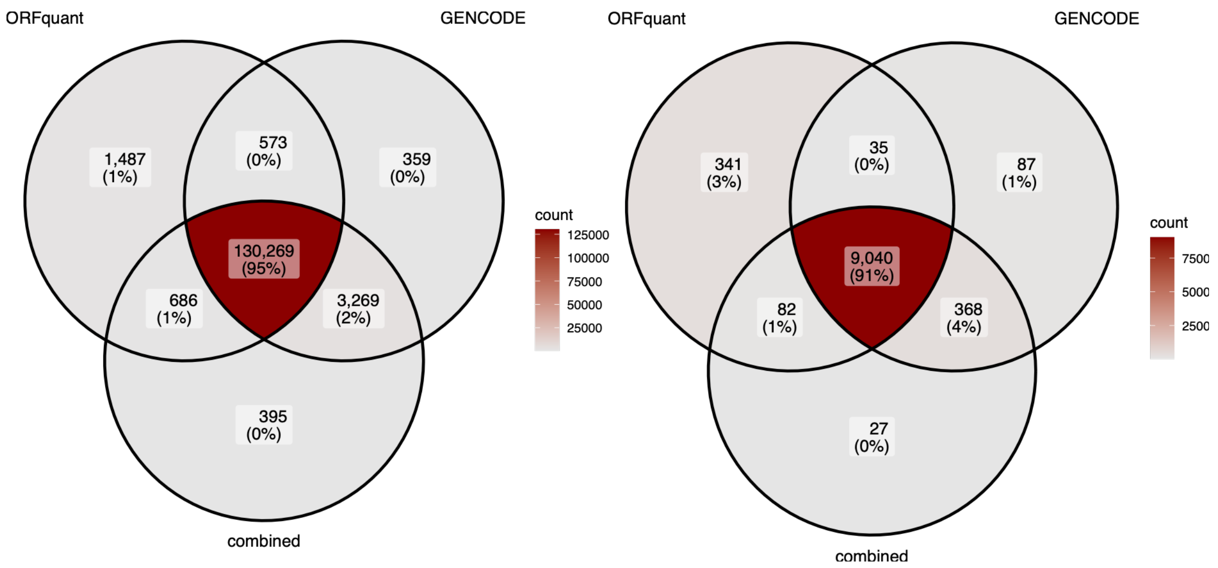


**Supplementary Figure 8** – Peptide identification across protein databases.

Venn diagram showing the number of peptides (left) or genes (right) detected in the TMT dataset by the different database searches.

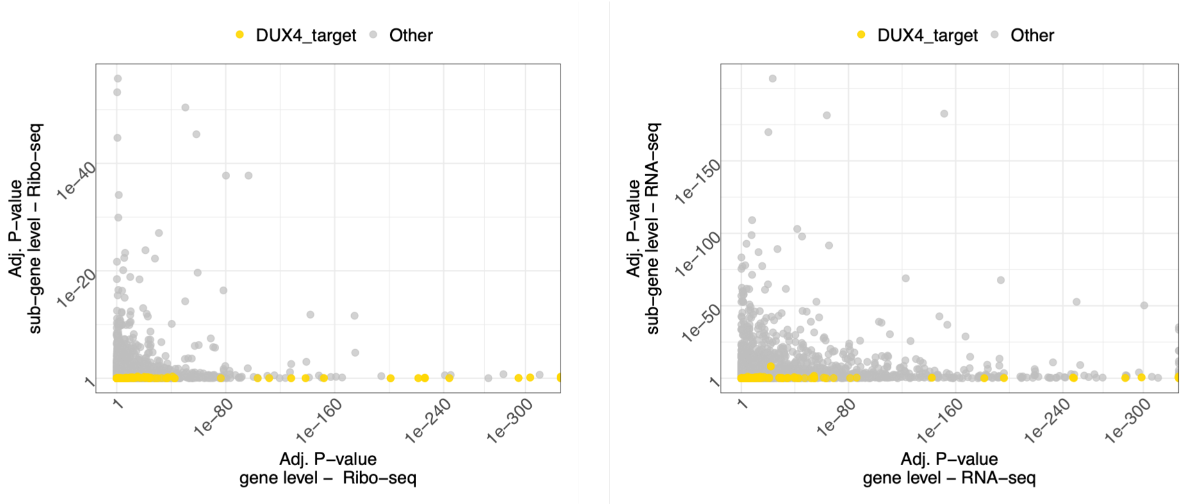


**Supplementary Figure 9** – Gene- and sub-gene- level regulation in Ribo-seq and RNA-seq.

P-values from gene-level analysis using *DESeq2* (y-axis) against P-values derived from transcript features (exonic bins and junctions) from *DEXseq* (x-axis), using RNA-seq or Ribo-seq.

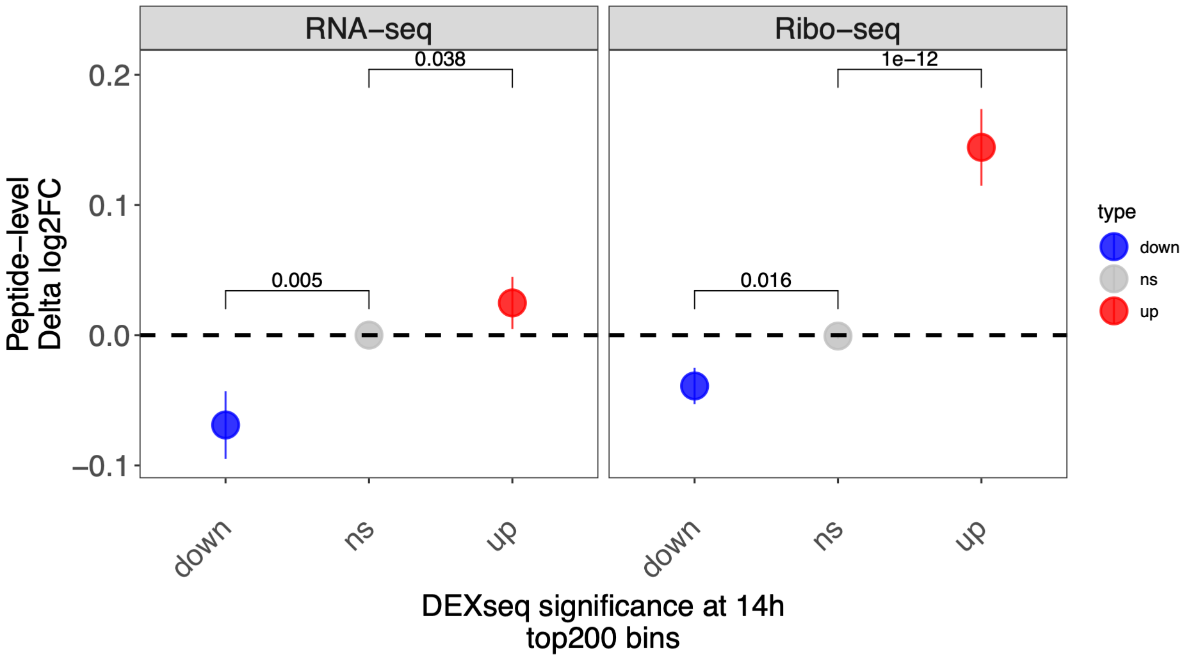


**Supplementary Figure 10** – Sub-gene agreement between RNA and protein using the top 200 DE bins.

Delta log2FC of peptide abundances divided according to the regulation of their originating exonic bin, using RNA-seq or Ribo-seq-derived significant transcript features (top 200 bins each). P-values from paired Wilcoxon test against values from peptides coming from non-changing exonic bins are shown.

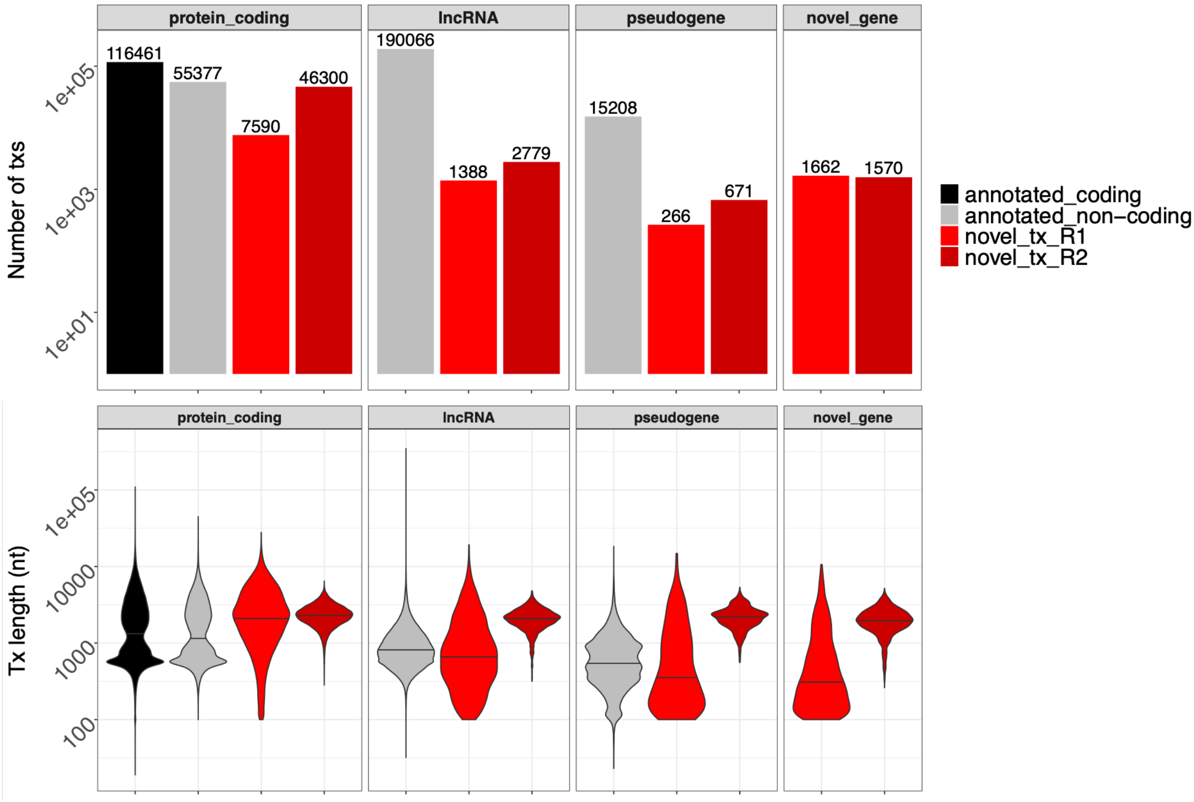


**Supplementary Figure 11** – Novel transcripts across annotations and gene biotypes.

Number of annotated, novel StringTie-assembled, and long-read-derived transcripts are shown above, with their lengths in nucleotides in the bottom section.

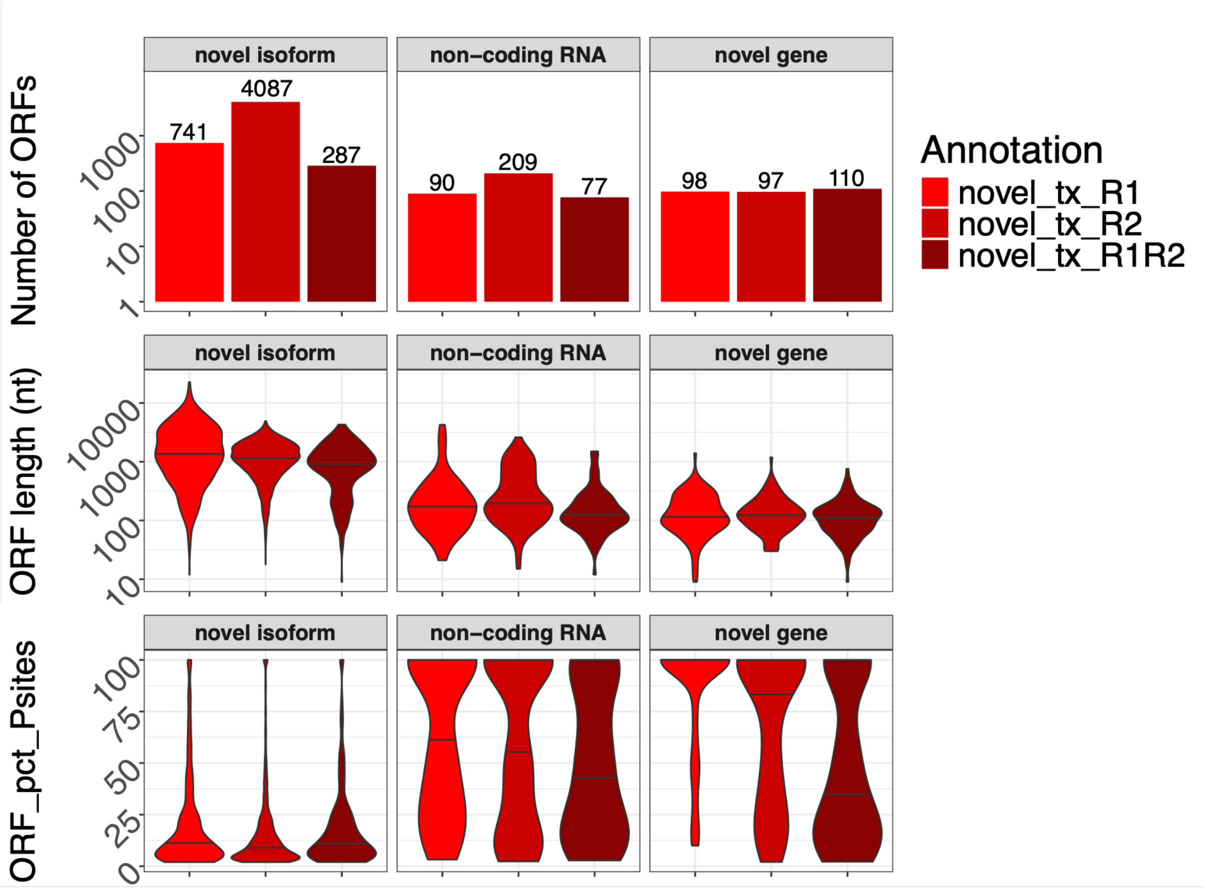


**Supplementary Figure 12** – Novel detected ORFs across annotations and gene biotypes.

Number of annotated, novel StringTie-assembled, and long-read-derived transcripts are shown above, with their lengths in nucleotides in the bottom section.

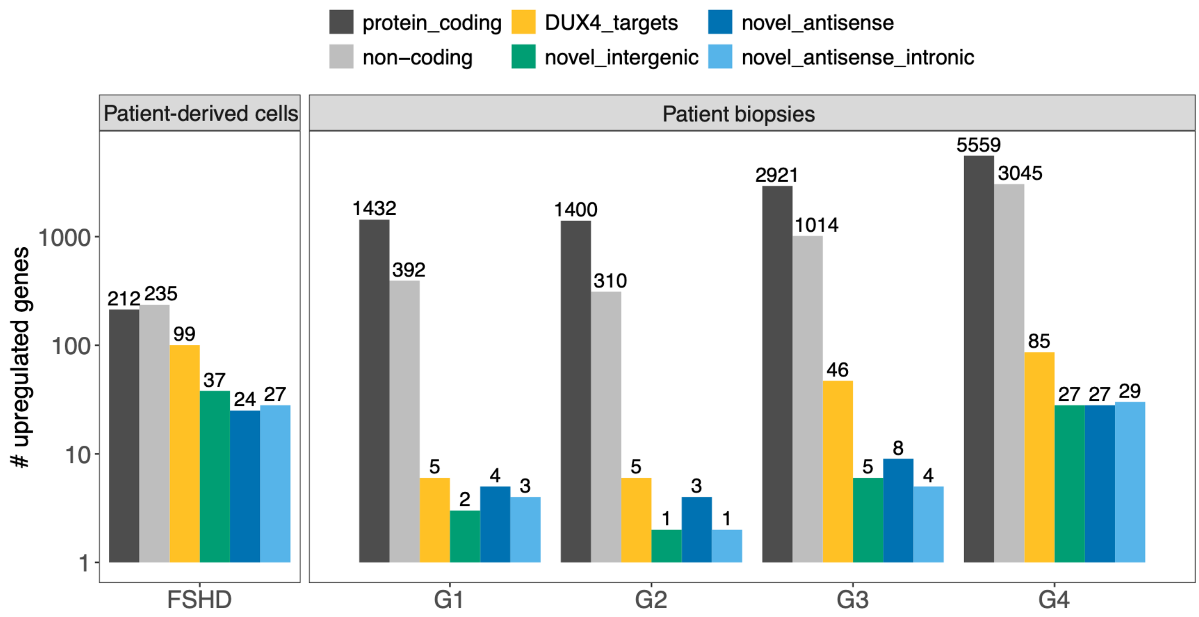


**Supplementary Figure 13** – Gene-level differential analysis in FSHD.

Number of upregulated genes in Yao *et al*, and Wang *et al* datasets, divided by different gene biotypes
